# Reconciling contrast invariance and non-linear computation in cortical circuits

**DOI:** 10.1101/2021.04.23.441165

**Authors:** L. Bernáez Timón, P. Ekelmans, S. Konrad, A. Nold, T. Tchumatchenko

**Affiliations:** Max Planck Institute for Brain Research, Theory of neural dynamics group; University of Bonn, Institute of experimental epileptology and cognition research; Frankfurt Institute for Advanced Studies, Johann Wolfgang Goethe University; Heidelberg University, Institute for Theoretical Physics

## Abstract

Network selectivity for orientation is invariant to changes in the stimulus contrast in the primary visual cortex. Similarly, the selectivity for odor identity is invariant to changes in odorant concentration in the piriform cortex. Interestingly, invariant network selectivity appears robust to local changes in synaptic strength induced by synaptic plasticity, even though: i) synaptic plasticity can potentiate or depress connections between neurons in a feature-dependent manner, and ii) in networks with balanced excitation and inhibition, synaptic plasticity is a determinant for the network non-linearity. In this study, we investigate whether network contrast invariance is consistent with a variety of synaptic states and connectivities in balanced networks. By using mean-field models and spiking network simulations, we show how the synaptic state controls the non-linearity in the network response to contrast and how it can lead to the emergence of contrast-invariant or contrast-dependent selectivity. Different forms of synaptic plasticity sharpen or broaden the network selectivity, while others do not affect it. Our results explain how the physiology of individual synapses is linked to contrast-invariant selectivity at the network level.

## Introduction

Neurons in sensory cortical areas selectively respond to certain features. For instance, neurons in the primary visual cortex (V1), in the primary auditory cortex (A1) and in the piriform cortex are selective to the orientation of visual images [23], to the frequency of sounds [37], or to the odorants in a scent [60], respectively. At the same time, cortical neurons respond to the stimulus intensity, which is measured as contrast in visual images [52], as sound pressure level in auditory cues [50], or as the concentration of odorant molecules in a scent [5]. In the piriform cortex, the response of selective neurons to both odorant and concentration has been studied [5]. Similarly, in V1, the response of orientation-selective neurons to orientation and contrast has been extensively characterized [23, 52, 4, 16]. But, one response mechanism that is still not yet fully understood neither in V1 nor in the piriform cortex is the emergence of contrast (or concentration) invariance at the network level [10, 8]. Specifically in V1, groups of neurons with a particular orientation preference show increased firing rates when a stimulus of a similar orientation appears in the environment, or when the contrast of a stimulus increases [10, 8] (Figure 1a-b). Interestingly, while the neuronal response to contrast is often non-linear, the response of the population to orientation is re-scaled by contrast, leaving its shape invariant [8] (Figure 1c,i). Computationally, contrast invariance can only emerge if the network activity is re-scaled by the same factor upon changes in contrast at all orientations within the network, regardless of the type of non-linear response to contrast [10]. However, if groups of neurons coding for certain orientations were more or less susceptible than others to the contrast of the input, the response to orientation could narrow (Figure 1c,ii) or broaden (Figure 1c,iii), which does not seem to be the case in V1 according to experimental reports [10, 8]. What mechanisms could give rise to orientation-independent re-scaling by contrast of the network response? Or vice versa, what mechanisms prevent it?

**Figure 1:**
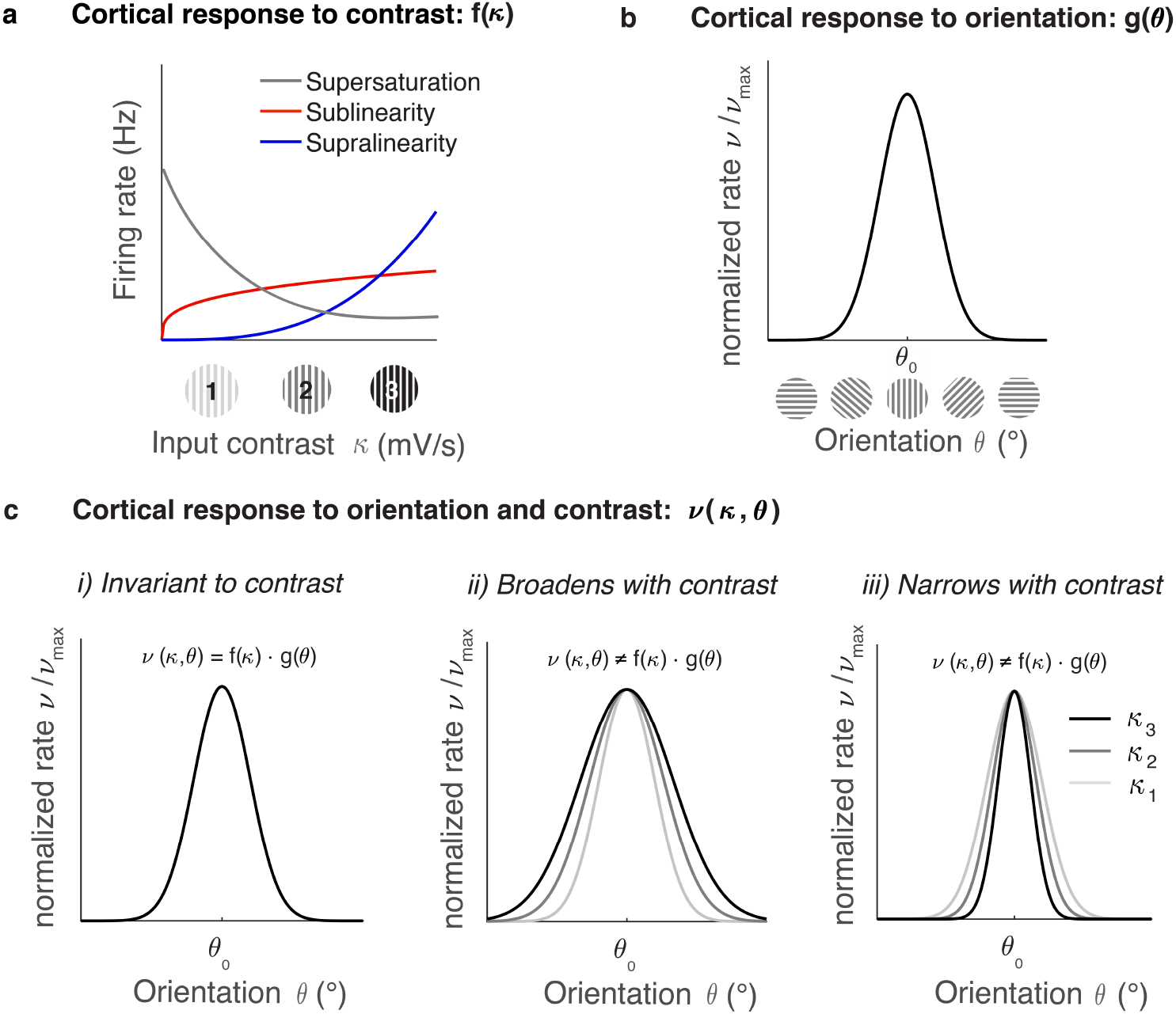
Possible outcomes for orientation and contrast in response to visual stimuli. **(a)** Visual stimuli of any orientation can appear at different contrasts in the environment depending on the illumination conditions (see *κ*_1_ < *κ*_2_ < *κ*_3_). Experiments report three types of non-linear cortical response to contrast *f* (*κ*): supersaturation, sublinearity, and supralinearity. **(b)** Neurons with similar orientation tuning in V1 are preferentially connected. Here, we illustrate a stereotypical response *g*(*θ*) of a cortical population to a stimulus of orientation *θ*_0_. The function *g*(*θ*) is also referred to as *tuning curve*. **(c)** Population response to the stimulus orientation and contrast *ν*(*κ*, *θ*) is a combination of the type of contrast response *f* (*κ*) depicted in (a) and the orientation tuning properties *g*(*θ*) shown in (b). Based on *f* (*κ*) and *g*(*θ*), three types of network response *ν*(*κ*, *θ*) are possible. **(i)** If contrast re-scales evenly the firing rate by the same factor at all active orientation domains, *ν*(*κ*, *θ*) is invariant to contrast. On the contrary, *ν*(*κ*, *θ*) tuning becomes contrast-dependent if the firing rates to different orientations are unevenly rescaled by contrast: tuning width broadens with contrast **(ii)** if the non-preferred orientations are more susceptible to increasing contrast or it narrows with contrast **(iii)** if the neurons with preferred orientation are more susceptible to increasing contrast.

Let us briefly review studies on invariant representations. Historically, contrast invariance was first reported in single neurons in V1 [52, 4, 16, 17]. A combination of neuronal mechanisms including gain-control [17], contrast-dependent noise [17, 4, 9] or a power-law input-output neuronal transfer function [19, 35, 42] were proposed to explain the emergence of contrast invariance in single neurons. The aforementioned mechanisms shape the transfer function of single neurons such that their output is contrast-invariant. Interestingly, a balance between excitation and inhibition has been reported in somatosensory cortices [39, 21] where network contrast invariance is present. Theoretical studies have shown that in E/I balance, the neuronal transfer function does not influence the activity of the network [58, 46], because its current contribution cancels in the limit of large networks. Therefore, contrast invariance at the network level cannot necessarily be explained as the sum of the activity of contrast-invariant neurons and needs to be studied at the network level.

In a network, the activity of V1 neurons is driven by the thalamic input, which is tuned for orientation [15, 13] and whose magnitude increases with contrast [12]. The activity of V1 neurons in response to the thalamic input spreads through the network via intracortical connections, which are established preferentially, but not exclusively, between neurons of similar orientation preference [27]. In V1, the strength of intracortical connections changes due to short-term synaptic plasticity (STP), which is ubiquitous in the synapses of pyramidal neurons [59]. In STP, changes in presynaptic firing rate affect the neurotransmitter release probability and induce the amplification or depression of thalamic input in a time-scale of milliseconds to seconds [1, 56]. This modifies the activity of the network and can compromise the emergence of contrast invariance [38]. Similarly, long-term changes in the synaptic strength can modify selectivity, both at the single neuron level [48] and at the network level [18]. What are the mechanisms enabling V1 to exhibit robust contrast invariance?

Here, we investigate candidate mechanisms giving rise to contrast invariance in balanced cortical circuits with plastic synapses. In particular, we look for transformations that simultaneously permit two features: i) the re-scaling of the network response by a constant at all features, and ii) the non-linear changes in the contrast response function induced by plasticity. To this end, we examine rate models in the limit of excitation-inhibition balance and test their predictions using spiking networks of leaky integrate-and-fire neurons. Our results reveal how STP generally leads to contrast-dependence. We also report a synaptic plasticity rule that reconciles both the emergence of contrast invariance and non-linear contrast response.

## Results

In this study, we examine how synaptic plasticity influences network response to contrast and how it shapes orientation selectivity. The dependence of firing rate on stimulus contrast (i.e. intensity) is known as the contrast response (Figure 1a). Neurons in cortical networks exhibit a variety of non-linear contrast response functions [3, 11]. For instance, the dependence of firing rate on the stimulus contrast can be amplified (Figure 1a, supralinearity) or reduced (Figure 1a, sublinearity). It is also possible that the firing rate decreases as a function of increasing contrast (Figure 1a, supersaturation). In addition, the network response to a stimulus of a given orientation is defined as the network selectivity for orientation (Figure 1b). The selectivity for orientation depends on the stimulus contrast (Figure 1c). A network is contrast-invariant if contrast does not modify its selectivity for orientation (Figure 1c,i). Conversely, a network is contrast-dependent if contrast broadens or narrows its selectivity for orientation (Figure 1c,ii-iii).

We study the interplay between synaptic plasticity, the contrast response and the selectivity for orientation using rate models and spiking network simulations (see Methods for details). For this purpose, we examine four types of synaptic interactions: constant, short-term facilitating, short-term depressing [36], and power-law synapses (see Methods). The spiking network models that we examined consist of leaky integrate-and-fire (LIF) excitatory (E) and inhibitory (I) neurons.

### How synaptic plasticity modulates the network response to contrast

Theoretical studies suggested that the non-linearities in the contrast response (Figure 1a) can be explained through synaptic plasticity in balanced networks [20, 36]. Here we examine which type of synaptic plasticity enables a particular response to contrast in balanced networks. To that end, we introduce the susceptibility to input contrast *δ* as an index. This is a measure for the relative change in the excitatory firing rate *ν_E_* as a function of the input contrast *κ*,

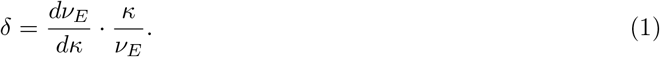

*δ* defines regimes of four types of contrast response functions (Figure 2a): supersaturation (*δ* < 0), sublinearity (0 < *δ* < 1), linearity (*δ* = 1), and supralinearity (*δ* > 1). We assume excitatory input contrasts (*κ* > 0).

**Figure 2:**
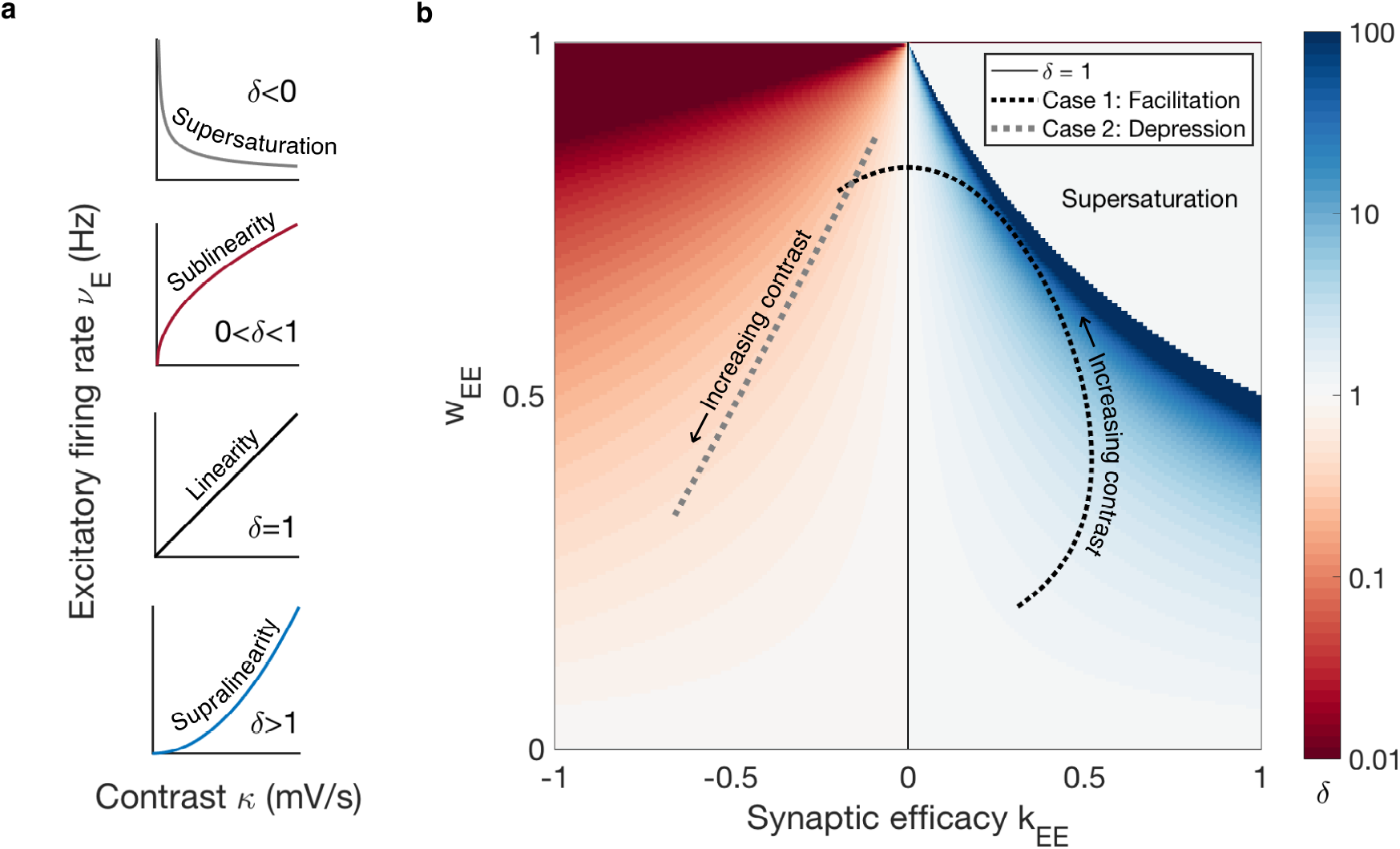
Plastic *E* → *E* synapses control the susceptibility to input contrast *δ* in balanced networks. **(a)** The contrast response characterized by the value of *δ*: supersaturating (*δ* < 0), sublinear (0 < *δ* < 1), linear (*δ* =1), and supralinear (*δ* > 1). **(b)** The phase space of values for *δ* (Equation 2) reveals a sublinear contrast response for depressing states (*κ_EE_* < 0), and a supralinear or supersaturating response for facilitating states (*κ_EE_* > 0). The network response is linear if synapses are constant (*κ_EE_* = 0, black line). The transition from the sublinear to the supralinear region is smooth, whereas the transition from the supralinear to the supersaturating is discontinuous and it is characterized by *δ* → ∞. The sublinear and supralinear non-linearities are stable and can be dynamically reached by the network, whereas supersaturation is unstable and may not be realizable in balanced networks (see Appendix). The trajectories describe how *δ* evolves as a function of increasing input contrast in a network with facilitating STP synapses (Case 1) and in a network with depressing STP synapses (Case 2) (see Methods - Model II and Table S1 for parameters). Let us note that for positive firing rates and a stable balanced state, parameters must satisfy *w_EE_* < 1 [58, 46].

In a balanced E/I network with synaptic plasticity in the *E* → *E* connections (Equation 10), *δ* can be elegantly written as (see Methods for derivation)

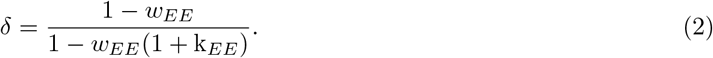

Here, *w_EE_* denotes the dimensionless synaptic *E* → *E* strength for a particular firing rate, 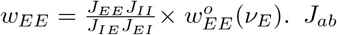 are the constant weights for synapses connecting population *b* to population *a*, where *a,b* ∈ {*E,I*}, and 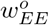 is the firing-rate dependent scaling factor of the *E* → *E* synapses. 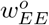 can also be interpreted as the probability of neurotransmitter release in STP. k_*EE*_ indicates how the synaptic strength changes with the firing rate:

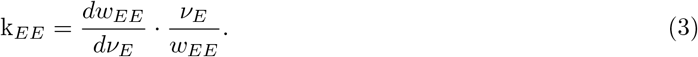

The synaptic state is depressing if k_*EE*_ < 0 and facilitating if k_*EE*_ > 0. The phase space of values for *δ* (Figure 2b) shows that depressing states (k_*EE*_ < 0) lead to sublinear contrast response. Moreover, we find that supralinearity and supersaturation are only possible for facilitating synaptic states (k_*EE*_ > 0). This is illustrated by the trajectories of the network response to increasing input contrast for networks with depressing and facilitating STP (Equation S3). The network with depressing synapses describes a trajectory in the sublinear space. The network with facilitating synapses occupies the supralinear space for positive values of k_*EE*_ while it transitions smoothly to the sublinear space as *w_EE_* saturates. A linear response only occurs if synapses are constant (k_*EE*_ = 0). Additional *E* → *I* plasticity also permits the emergence of all types of non-linearities (Figure S2, Equation S6).

There are two possible transitions between regimes. One is a smooth transition from the sublinear to the supralinear region (vertical black line at k_*EE*_ = 0). In this case, *δ* varies continuously from values smaller than unity to values larger than unity. The other is a discontinuous transition from the supralinear regime to the supersaturating regime. This discontinuous transition is characterized by diverging network susceptibility *δ* → ∞. If a network approaches this transition from the supralinear regime, it will experience a rapid increase in firing rate. Facilitating short-term synapses cannot support this rapid increase in firing rate, leading to a transition of the trajectory to the sublinear regime. Network states in which *δ* < 0 are unstable (see Appendix). This result indicates that sublinearity, linearity, and supralinearity are realizable in balanced networks through plastic synapses. Supersaturation, however, is not supported in balanced networks with synaptic plasticity.

In the next section we analyze the network response in a spiking network with STP.

### Susceptibility *δ* in spiking networks with STP

In this section we study if the different types of contrast response functions reported in Figure 2 emerge in a spiking network with *E* → *E* short-term synaptic plasticity (Equation S3, Figure S3). STP modulates the synaptic strength as a function of the firing rate history of the presynaptic neuron [55], and acts on the timescale of milliseconds to seconds.

We examine the response to contrast in the spiking network model illustrated in Figure 3 (see Model II in Methods). Spiking networks with STP facilitating and depressing *E* → *E* synapses have a non-linear response to contrast (Figure 3b, dots). The synaptic scaling factor 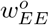 and the synaptic efficacy k_*EE*_ in these networks change with the firing rate (Figure 3d-e, dots). In contrast, the network with constant synaptic weights is linear (Figures 3b, black dots, and Figure S1). We confirm that these networks behave in agreement with the analytical balanced state prediction (Equation 10, Figure 3, solid lines). The CV ≈ 1 values and exponentially decaying ISI distributions (Figure 3c) are consistent with in vivo recordings [53], and demonstrate that STP does not modify the irregularity in spiking activity (see Figure S1c-d for reference).

**Figure 3:**
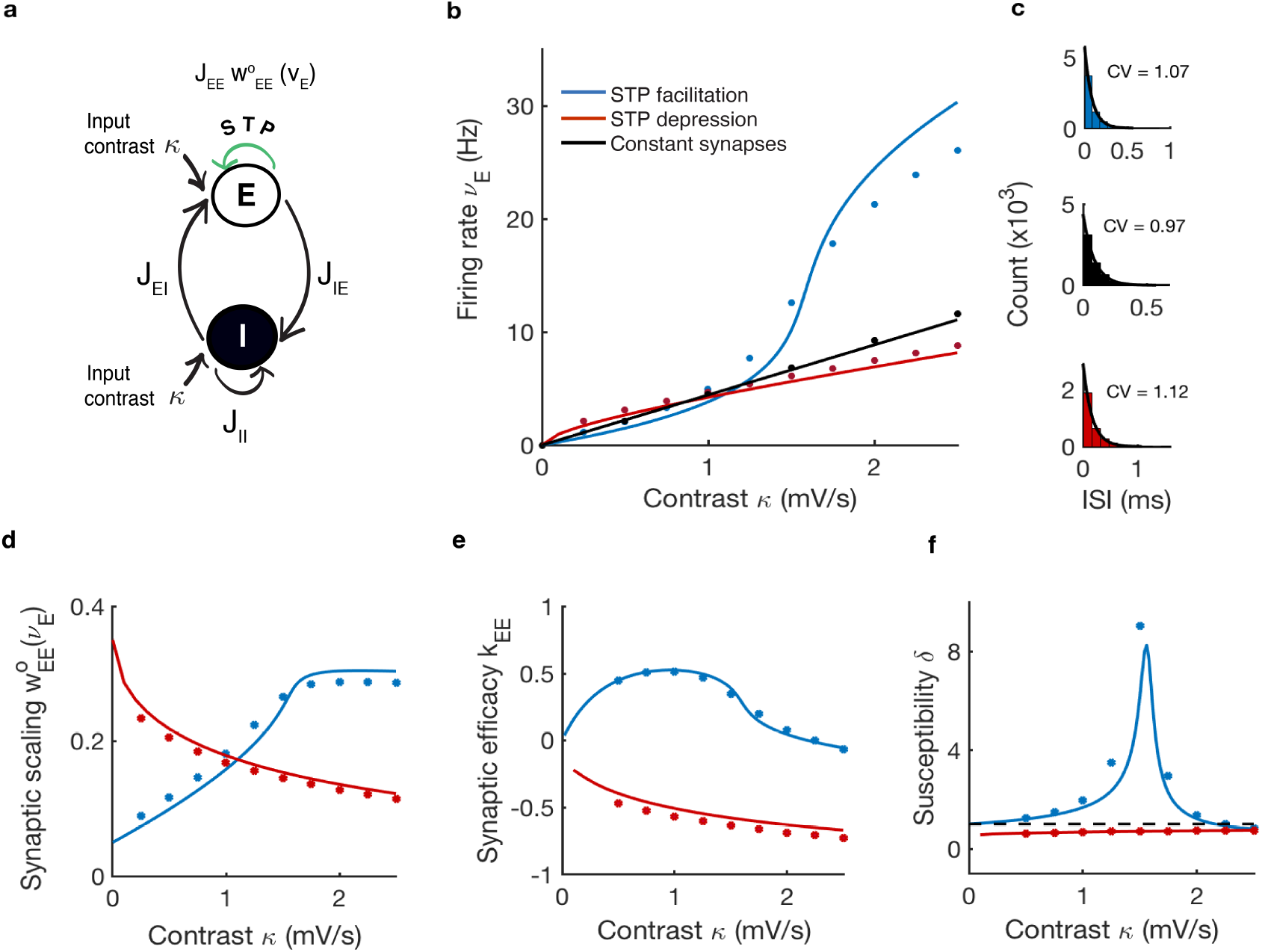
Susceptibility to contrast *δ* in spiking networks of randomly connected neurons with STP in the *E* → *E* synapses. **(a)** Homogeneously randomly connected network (see Methods - Model II and Table S1 for parameters). The *E* → *E* synaptic strength is modulated by the function 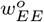, which describes the probability of neurotransmitter release as a function of the presynaptic firing rate (green arrow). Each neuron receives an external excitatory input that is directly proportional to the stimulus contrast *κ* > 0. **(b)** The firing rate of the *E* population vs the input contrast for constant, facilitating and depressing STP synapses. Data from spiking network simulations (dots) is shown alongside balanced network predictions from Equation 10 (solid lines). **(c)** The coefficient of variation CV≈ 1, ISI histograms and exponential fit denote poissonian irregular firing patterns. Data computed over 1 second of spiking network simulation for an input contrast of 1.5 mV/s. **(d)** The probability of neurotransmitter release 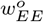 and **(e)** the synaptic efficacy k_*EE*_ in the two plastic networks studied. Predictions from the rate model (solid lines) are plotted alongside data from spiking network simulations (dots). **(f)** The susceptibility *δ* calculated from Equation 2 (solid lines) and its value in spiking network simulations (dots). The dashed line indicates *δ* = 1.

We calculate *δ* in the spiking networks based on the values of 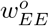 and k_*EE*_ displayed by plastic synapses (Figure 3f). As predicted, the spiking network with facilitating synapses exhibits a supralinear increase in firing rate (*δ* > 1) that becomes sublinear (0 < *δ* < 1) for increasing contrasts (Figure 3f, blue). This behavior is consistent with the facilitating *δ*-trajectory in Figure 2. For low firing rates, neurotransmitter is replenished fast enough in between spikes and facilitating synaptic transmission is sustained. The value of *δ* increases and the trajectory approaches the supersaturating unstable regime, where *δ* → ∞. This explains the peak in *δ* in Figure 3f. At this point, a further increase in synaptic strength can not be physiologically supported because synapses demand neurotransmitter quicker than it can be produced. As a result, the network transitions smoothly from the supralinear to the sublinear regime. Conversely, the network with depressing synapses exhibits a sublinear increase in firing rate (0 < *δ* < 1) for all the input contrasts analyzed, which is also in agreement with the trajectory in Figure 2.

We show that short-term synaptic plasticity can have a dramatic effect on the non-linear contrast response in spiking networks. In addition, we demonstrate that *δ* can be used as a tool to study the behavior of spiking networks with synaptic plasticity. Next, we apply these tools to networks with orientation selectivity.

### Networks with constant synaptic weights are contrast-invariant and have a linear contrast response

In this section, we run a ‘‘control experiment” in which we examine the response to contrast and orientation selectivity in networks with constant synapses.

For this, we model networks of neurons that are selective for the orientation of the stimulus (Figure 4a). In the model, neurons acquire their orientation preference from a Gaussian stimulus representing thalamocortical input (see Figure S6). The preferred orientation *θ*_0_ is set by the location of the peak of the Gaussian. The stimulus contrast is a pre-factor *κ* that scales the magnitude of the thalamocortical input [16]. In addition, the probability of connection between neurons decays as a Gaussian with orientation-distance, such that neurons with a similar orientation preference are more likely to be connected [27, 29] (see Model III in Methods). In a network with constant synapses, these features yield the network selectivity for orientation *ν_E_* [46]:

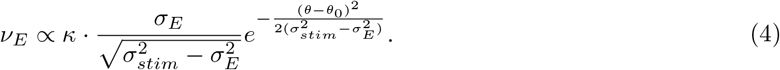

**Figure 4:**
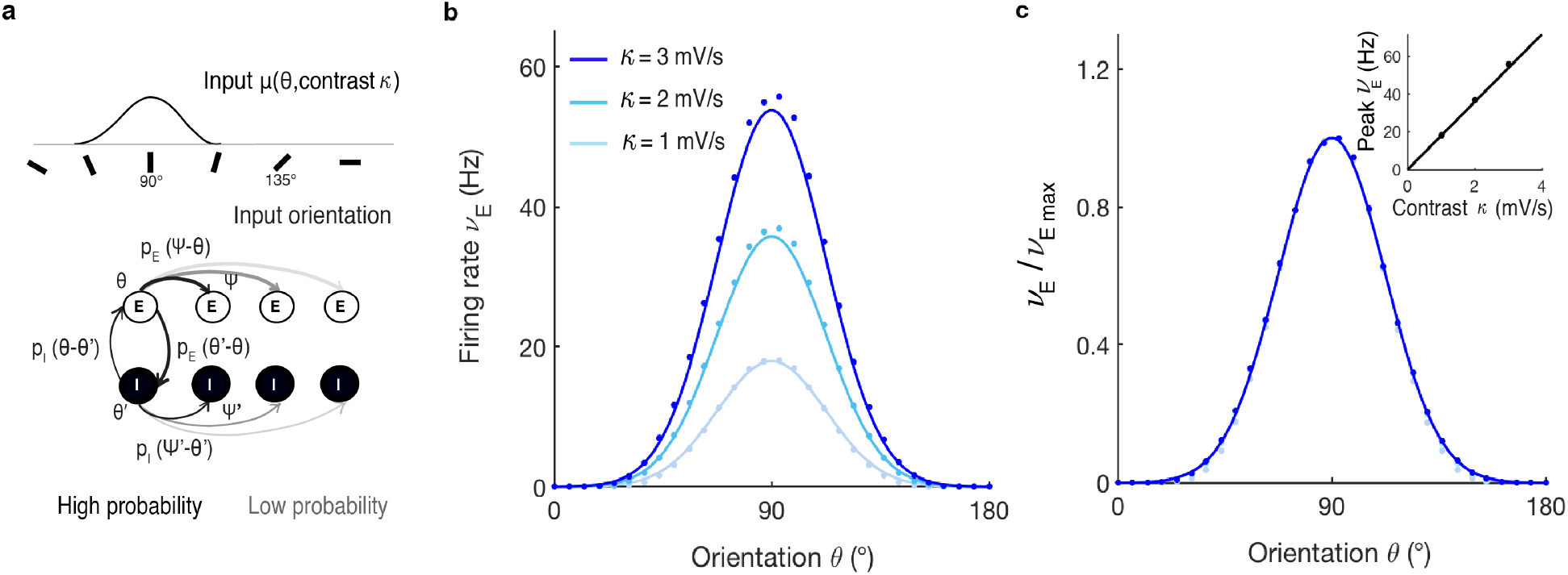
Networks with orientation-dependent connectivity and constant synapses have linear susceptibility and are contrast-invariant. **(a)** A Network with feature-dependent connectivity (see Methods - Model III and Table S1 for parameters). The connection probability functions *p_E_* and *p_I_* are Gaussian. **(b)** Excitatory tuning curves in response to different input contrast, *κ* = 1, 2, 3 (mV/s). Prediction from the balanced theory (Equation 4, solid lines) compared to the results obtained in a simulation of spiking neurons (dots). **(c)** Overlapping normalized tuning curves from (b) show contrast-invariance. Inset: peak firing rate at *θ* = *θ*_0_ from tuning curves in (b) linearly increases with contrast.

Here, *ν_E_* denotes firing rate as a function of orientation, *κ* is the contrast of the stimulus, *σ_stim_* is the width of the thalamic input, and *σ_E_* is the width of the connectivity from *E* → *E* and from *E* → *I* neurons.

Equation 4 and the spiking network results shown in Figure 4 demonstrate that orientation selectivity is Gaussian in balanced networks with constant synapses. These results also show that the stimulus contrast *κ* linearly re-scales firing rates at all orientations without changing the shape of *ν_E_* (Figure 4b-c, solid lines). The factorization of *ν_E_* (Equation 4) into a term that depends only on contrast and a function that depends only on orientation *θ* implies that the network is contrast-invariant. This is shown in Figure 4c, where we normalize *ν_E_* by the respective peak firing rates shown in Figure 4b. The overlap of all orientation selectivity curves confirms that networks with constant synapses are contrast-invariant. The contrast response of the network is linear and *δ* = 1 (Figure 4c, inset for *θ* = *θ*_0_).

This ‘control experiment” demonstrates that networks with constant synapses have a linear response to contrast and contrast-invariant orientation selectivity. Can networks with synaptic plasticity that exhibit sublinear or supralinear susceptibility also be contrast-invariant?

### STP yields contrast-dependent orientation selectivity

In this section, we study if a non-linear contrast response induced by short-term synaptic plasticity is compatible with contrast invariance in networks with orientation-dependent connectivity (see Model IV in Methods, Figure 5a).

**Figure 5:**
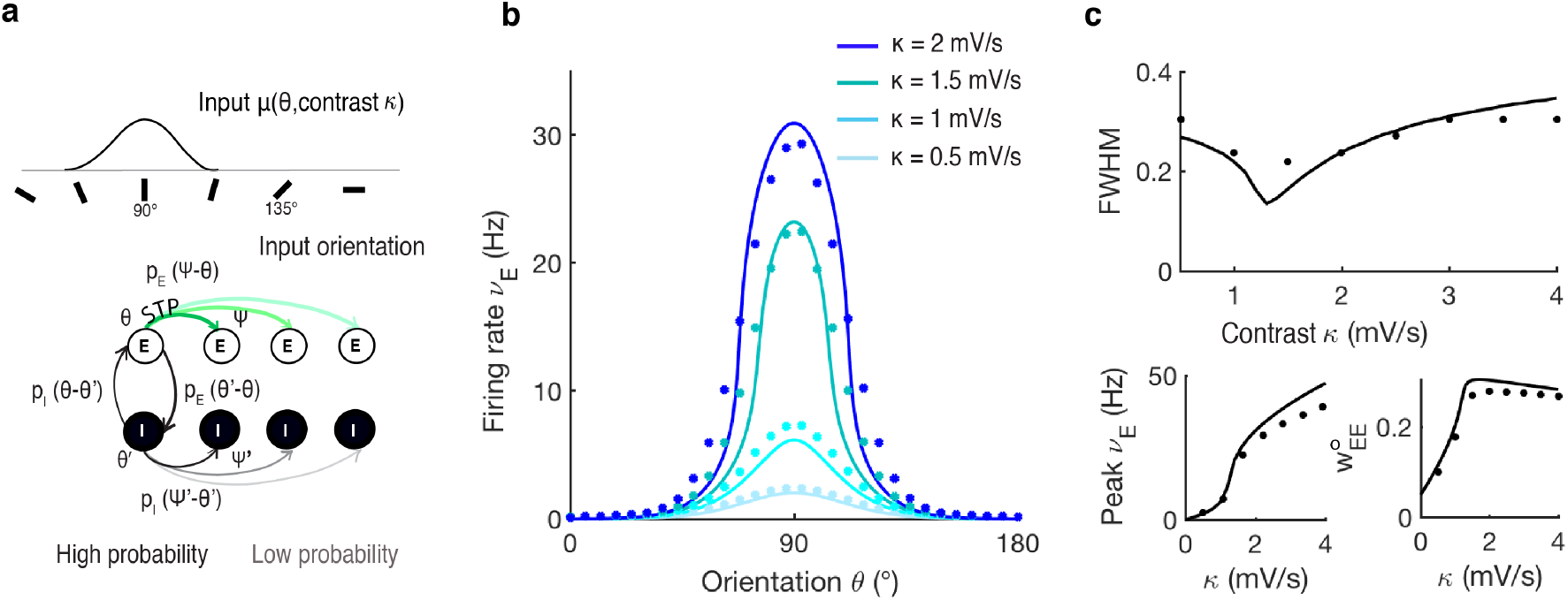
STP in the *E* → *E* synapses yields non-linear susceptibility to input contrast and makes networks contrast-dependent. **(a)** Network with orientation-dependent connectivity and *E* → *E* STP synapses (see Methods - Model IV and Table S1 for parameters). **(b)** Excitatory tuning curves for different input contrast, *κ* = 0.5,1,1.5, 2 (mV/s). Predictions from the balanced theory (Equation 5, solid lines) compared to the results obtained in a simulation of spiking neurons (dots). **(c)** Upper pannel: the full width at half maximum (FWHM) changes with contrast, which confirms contrast-dependent selectivity. Lower pannel: the peak firing rate increases supralinearly until *κ* approaches 1.5 mV/s (left), which is consistent with the increase in the probability of neurotransmitter release 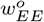 (right). As a signature of facilitating transmission, when firing rates continue to increase for *κ* > 1.5 mV/s, the probability of neurotransmitter release 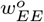 decreases (right) and the network behavior becomes sublinear (left). The shif from supralinearity to sublinearity is consistent with the trajectory in Figure 2 (Case 1).

Through the susceptibility index *δ*, we have shown how *E* → *E* STP controls the network response to contrast in balanced networks of randomly connected neurons (Figures 2-3). Here, we show that STP also controls the non-linearity in the response to contrast in networks with orientation-dependent connectivity. In this type of network, the response to contrast and the selectivity for orientation are related through (see Appendix)

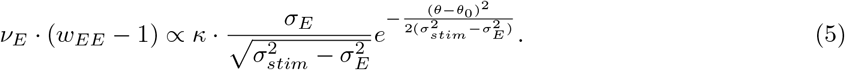

Note that here, *w_EE_* is proportional to the probability of neurotransmitter release in STP. In Figure 5, we show results for facilitating synapses (for depressing synapses see Figure S4).

The solution of Equation 5 is given in Figure 5b (solid lines) for several input contrasts *κ*. When synapses are facilitating, increasing the stimulus contrast *κ* does not linearly re-scale *ν_E_*. This effect is also present in spiking networks (Figure 5b, dots). We quantify the contrast dependence on selectivity using the full width at half maximum (FWHM) (Figure 5c). With an increase in contrast, the FWHM first decreases, indicating a narrowing. Interestingly, after reaching a minimum, the FWHM then increases, revealing a subsequent broadening. These results show that networks with STP in the *E* → *E* synapses are contrast-dependent. This dependency on contrast is consistent with the facilitating trajectory in Figure 2. For a given input contrast, the neurons tuned to the stimulus orientation *θ*_0_ receive higher input than any other position in the network, and the firing rate increases. As a result, the point of maximum supralinearity is reached faster and the value of *δ* is higher than the one at positions that have a different preferred orientation, which narrows the selectivity. If at this point the contrast increases, synapses at the position tuned to the stimulus orientation can no longer meet the demand for neurotransmitter required to sustain high firing rates, which causes a decrease in *δ*. At the same time that the position tuned to the stimulus orientation approaches sublinearity, positions tuned for the non-preferred orientations approach the point of maximum supralinearity, which broadens the selectivity (see Video S1). The transition between non-linear *δ* regimes is also seen in the network response to contrast and in the STP probability of neurotransmitter release (Figure 5c, for *θ* = *θ*_0_). These results demonstrate that the non-linear contrast response induced by STP leads to contrast-dependent orientation selectivity.

### Power-law synapses support non-linear contrast response and contrast-invariant selectivity

In this section, we show that the reconciliation between the non-linear contrast response and the emergence of contrast invariance is plausible under certain synaptic conditions. Previous theoretical studies have reported that only a power-law function can transform contrast-invariant membrane potential input into contrast-invariant firing rate output in single-neurons [19, 35]. Here, we impose a power-law transformation at the synaptic level at *E* → *E* and *E* → *I* synapses, 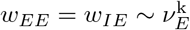 (see Model V in Methods). Note that both a power-law plasticity in the *E* → *E* (Equation 5) and STP in the *E* → *E* and *E* → *I* connections (Figure S5) lead to contrast-dependence. Instead, here the response to contrast and the selectivity for orientation are related through (see Appendix)

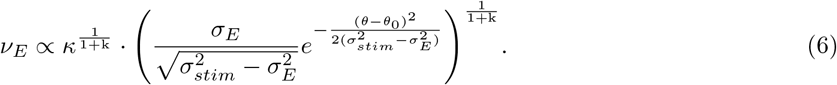

Here, the synaptic efficacy k is independent of the presynaptic firing rate. In this type of network, the susceptibility *δ* is given by 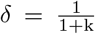. The network response is sublinear if k > 0, linear if k = 0, and supralinear if —1 ≤ k < 0.

The analytical solution of Equation 6 for k = —0.5 is given in Figure 6b (solid lines) for several input contrasts *κ*. Figure 6b shows that the activity of a spiking network with orientation-dependent connectivity and power-law synapses can be predicted by Equation 6. It is possible to factorize Equation 6 into a function of contrast and a function of orientation, which indicates contrast invariance. The normalization of the network activity demonstrates that contrast invariance is also present in spiking networks (Figure 6c, dots). As predicted, the contrast response is supralinear with *δ* = 2 (Figure 6c, inset for *θ* = *θ*_0_). This demonstrates that depressing synaptic states can lead to a supralinear response to contrast in spiking networks (Figure 6c). These results confirm that a power-law at the synaptic level is consistent with contrast invariance and a non-linear response to contrast at the network level.

**Figure 6:**
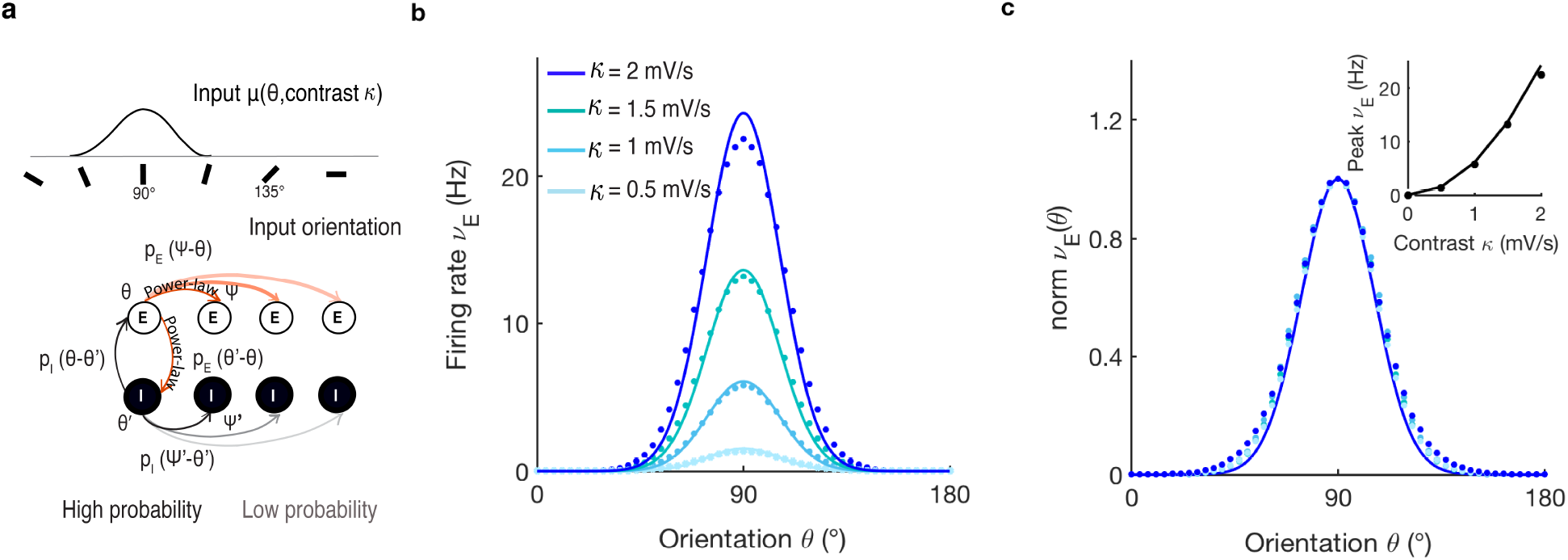
Networks with orientation-dependent connectivity and power-law *E* → *E* and *E* → *I* synapses show nonlinear susceptibility and contrast-invariant tuning. **(a)** Network with orientation-dependent connectivity. Additionally, in this network, the *E* → *E* and the *E* → *I* synapses are nonlinearly modulated by the function 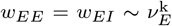, where k is the synaptic efficacy (red arrows) (see Methods - Model V and Table S1 for parameters). **(b)**. Excitatory tuning curves in response to several input contrast *κ* = 0.5,1, 1.5, 2 (mV/s). Predictions from the analytical balanced state (Equation 6, solid lines) are compared to the results obtained in a simulation of spiking neurons (dots). **(c)** Normalized analytical (solid lines) and simulation tuning curves (dots) from (b) show that this type of network is contrast-invariant. Inset: the peak excitatory firing rate at *θ* = *θ*_0_ increases supralinearly with contrast in the analytical balanced state (solid line) as well as in spiking network simulations (dots).

## Discussion

Here, we analyzed the interplay between synaptic plasticity, network response to stimulus contrast, and selectivity for the stimulus orientation. Counter-intuitively, contrast invariance of the network does not necessarily follow from the contrast invariance of individual neurons in balanced networks. Here, we showed how for balanced networks the synaptic state controls the non-linearity in the network response to contrast (Figures 2, 3). Depending on the synaptic state, the network can exhibit a variety of activation functions, from sub-to supralinear. Our results show that synaptic plasticity at specific connections describes a large range of interactions between contrast and feature selectivity and leads to contrast-invariant (Figure 6) or contrast-dependent (Figure 5) network selectivity. Therefore, this proposes a mechanism to reconcile a non-linear contrast response and contrast invariance at the network level that is compatible with E/I balance.

### Network response to contrast

In this work, we propose a measure to describe the non-linear behaviour of the network activity and its dependence on its synaptic states: the susceptibility to contrast *δ* (Equation 2). The susceptibility *δ* captures the relative changes in the excitatory firing rate with contrast. In balanced networks with *E* → *E* plastic synapses, we show that the susceptibility to contrast *δ* depends only on two variables: the effective synaptic strength *w_EE_* and its relative dependence on firing rate *ν_E_*, which we characterize by k_*EE*_ (see Equation 3). Using the concept of susceptibility *δ*, we identified four types of contrast responses: supralinear, linear, sublinear and supersaturating (Figure 2). In addition, the phase space of *δ* values allows to connect the synaptic state to phase transitions of network behavior (Figure 2, trajectories). We show trajectories spanning the broad phase space of non-linear responses for spiking networks with facilitating and depressing STP (Figure 3). We also show that a supersaturating non-linearity is unstable in balanced networks with plastic *E* → *E* synapses (Appendix).

These results apply to networks with E/I balance. Previous theoretical studies have shown that nonneuronal mechanisms like synaptic plasticity are necessary to achieve a non-linear response in balanced networks [36]. On the experimental side, studies have reported the cancellation of E/I currents in cortices where contrast invariance is present [39]. Furthermore, E/I balance has been proposed as a necessary mechanism for the emergence of sharp orientation selectivity in neurons located at pinwheels in V1 [21]. For completeness let us note that when excitation and inhibition are loosely balanced, the neuronal non-linearity has been shown to shape the network response to contrast. This scenario is captured by the supralinear stabilized network model (SSN), which considers constant synapses [2, 47]. The SSN permits supersaturation in rate models [2, 47] and in spiking networks [49]. Here, to single out the synaptic effect on the network nonlinearities, we considered balanced E/I networks with plastic synapses.

### Contrast-dependent selectivity

When investigating orientation selectivity in networks with different types of synapses and contrast responses, we found that for constant synapses the response to contrast is linear and the network is contrast-invariant (Figure 4). This is a consequence of the linearity of the network for all orientations *θ*. However, when synapses follow STP plasticity (Equation S3), the response to contrast is non-linear, and the network becomes generally contrast-dependent (Figure 5).

Addressing the consequences of STP, we found that facilitating states can narrow the network selectivity, while depressing states can broaden it (Figures 5, S4). This change in selectivity can be explained as follows: a stimulus of orientation *θ*_0_ preferentially depolarizes some neurons in the network. As a consequence, the firing rate *ν_E_*(*θ*_0_) of these neurons increases. This increase activates synapses, which start to release neurotransmitter with probability 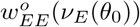. Conversely, neurons tuned to non-preferred orientations *ψ* hardly receive external input and *ν_E_* (*ψ*) < *ν_E_* (*θ*_0_), and their synapses release neurotransmitter with probability 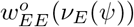. When the stimulus contrast increases, *ν_E_*(*θ*_0_) and *ν_E_*(*ψ*) increase, and the probability of neurotransmitter release changes at both orientations. The key to understanding this phenomenon is that in STP, the relative change in release probability depends on the starting firing rate value (Equation S3). Thus, the synaptic efficacy k_*EE*_(*θ*_0_) = k_*EE*_(*ψ*). As a result, if the synaptic state is facilitating, the firing rate at the network preferred orientation may be more supralinearly amplified compared to the orthogonal orientation. In this case, the network selectivity narrows. Conversely, if the synaptic state is depressing, the firing rate at the preferred orientation is dampened compared to the non-preferred orientations, which broadens the selectivity (see Video S1). Contrast-dependence has been observed in the auditory cortex, where the network selectivity for the frequency of a tone changes when measured at different sound intensity levels [44, 51, 30, 54, 37]. Therefore, our results suggest that STP may play an important role in modulating network selecitivity as a function of contrast.

### Contrast-invariant selectivity

The aforementioned scenario changes if the synaptic strength has a power-law like dependence as a function of firing rate. In this scenario, we found that a non-linear response to contrast is consistent with contrast invariance (Figure 6). Our results do not require that the synaptic strength of each synaptic terminal has to scale as a power-law function of the presynaptic firing rate, which would be unlikely due to the heterogeneity observed across synaptic terminals [7]. Instead, it is the average synaptic strength of a given population that should scale as a power-law for a circuit to be contrast-invariant. This condition must be fulfilled, at least, for excitatory synapses, regardless of whether the postsynaptic neuron is excitatory or inhibitory. Why do power-law synapses permit a non-linear response to contrast and contrast-invariant selectivity, while other synaptic plasticity rules (e.g. STP) do not? Previous studies suggest that a power-law is the only function that can transform contrast-invariant input into contrast-invariant spike output [35]. This is because powerlaw functions are scale-free transformations. In other words, a power-law synapse will amplify or compress the presynaptic rate by the same relative amount, independently of the firing rate of the presynaptic neuron. This ensures that the susceptibility *δ* is the same across orientations and for all firing rates. This universal non-linear re-scaling of the network activity supports contrast-invariant selectivity and permits the non-linearity in the response to contrast. Power-law transformations have been previously reported in the cortex at the neuronal level [17, 41]. Experiments showed that this type of transformation in single neurons can result from neuronal noise, which smooths the linear-threshold neuronal transfer function into a powerlaw [17, 41]. In agreement with this, theoretical studies demonstrate that a power-law transformation of membrane potential input into firing rate output is a requirement for single-neuron contrast invariance [19, 35]. However, a power-law neuronal non-linearity is insufficient to explain contrast invariance at the circuit level [8]. This is because in balanced networks the averaged activity is independent on the neuronal properties [58]. If that is the case and if a power-law transformation is necessary for the emergence of contrast invariance at the network level, then the power-law should be supported by synaptic, rather than by neuronal mechanisms in balanced networks. It is possible that neuronal and synaptic power-law non-linearities could coexist, each of them serving the function of contrast invariance at a different level. For example, contrast-invariant selectivity in single neurons and at the circuit level is found in the visual cortex [9, 8] and the piriform cortex [5]. In the piriform cortex, the selectivity for a particular odorant is not modified by the concentration of odorant molecules in the air. In addition, our results suggest that the emergence of contrast invariance or contrast dependence in cortical circuits may not require major anatomical differences. Indeed, numerous experiments report a similar columnar structure, the same number of cortical layers, similar cell types and similar input-output organization across sensory cortices [14, 45]. Instead, our results suggest that these differences in selectivity across auditory, visual and piriform cortices could be explained through synaptic physiology.

### Experimental validation

Our results suggest that differences between contrast-invariant cortices (i.e. visual, piriform) and contrast-dependent cortices (i.e. auditory) may result from constraints in the synaptic strength leading to experimentally testable predictions.

In particular, our results predict that excitatory synaptic strengths in contrast-invariant cortices should scale as a power-law with the excitatory presynaptic firing rate. But how can one assess the impact of the synaptic strength – which is a dynamic and distributed cellular property acting at the microscale – on the overall behavior of the network such as its firing rate? Given that the synaptic strength is sensitive to variations in the extracellular fluid, spontaneous activity, patterns of stimulation, and the presence of neuromodulators [6], it is important to study *in vivo* data. However, this is challenging. For example, one needs to identify cellular connections. One way could be to identify synaptically connected neurons guided either by single-cell optogenetic control of the presynaptic activity [40] or by fluorescent genetic labeling of specific cell types [24, 25]. Once synaptic strengths can be measured in a representative number of synaptic connections, differences in the synaptic physiology of invariant and non-invariant cortices can be studied, as our theoretical results suggest. In a next step, neuromodulators could be employed, which can change the function and dynamics of synapses and circuits [31], while the network activity could be recorded using two-photon calcium imaging. Indeed, the application of cholinergic agonists or antagonists has been shown to change the receptive field properties of single-neurons in the somatosensory [34] and auditory cortex *in vivo* [32]. As a neuromodulator, one promising candidate is adenosine: First, it down-regulates the release probability at excitatory synapses, while it has no or little effect on inhibitory transmission [26, 43].

Second, the mechanism mediating the decrease in release probability is presynaptic (as in our model): the activation of A1 adenosin receptors reduces the open probability of presynaptic Ca^2+^ channels [43]. Our results predict that contrast-invariant cortices loose this property when the synaptic behavior is disrupted by the administration of neuromodulators, while contrast-dependent cortices may experience changes in their selectivity properties.

### Conclusion

In summary, we have shown that different types of synaptic plasticity can generate a variety of non-linearities in the representation of input contrast in balanced networks. At the same time, synapses play a crucial role in the establishment of contrast-invariant or contrast-dependent selectivity. Our results indicate that the ability of cortical networks to extract invariant information about sensory stimuli is directly connected to the physiology of synapses.

## Methods

### Spiking network

We study five network models (I, II, III and IV and V). All the models consist of *N* neurons, of which *N_E_* = *q* · *N* are excitatory and *N_I_* = (1 – *q*) · *N* are inhibitory, with *q* = 0.8 [33]. Neurons are uniformly distributed in a one-dimensional state space of orientation preference. Let us remark that this defines the feature space, not the physical space. Neuron *i* in population *a* has the orientation preference 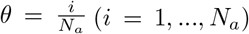, where *a* = {*E,I*}. Connected neurons with preferred orientation *θ* and *ψ* are sampled from a probability distribution ~ *p_a,b_*(*θ* – *ψ*). We consider a periodic domain Γ = [0,1], such that on 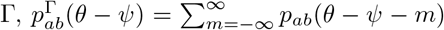. Let us note that in the figures we re-scale Γ to 180° for illustrative purposes. We assume *p_EE_* = *p_IE_* = *p_E_* and *p_EI_* = *p_II_* = *p_I_*. The membrane potential 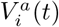 of neuron *i* from population *a* = {*E, I*} obeys the leaky-integrate-and-fire (LIF) dynamics described by

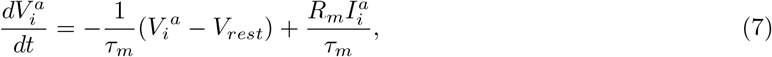

where *τ_m_* is the membrane time constant, *V_rest_* is the resting potential, *R_m_* is the membrane resistance, and 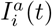 is the sum of the recurrent current from other neurons in the network and an external feed-forward current *F*_ffw_(*θ*). Whenever 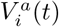 reaches the threshold voltage *V_th_* neuron *i* spikes and its membrane potential is reset to *V_rest_*. Each neuron in the network receives input from a fixed number of *C_E_* and *C_I_* presynaptic excitatory and inhibitory neurons, respectively. The total synaptic input to the *i*-th neuron from population *a* = {*E, I*} is given by

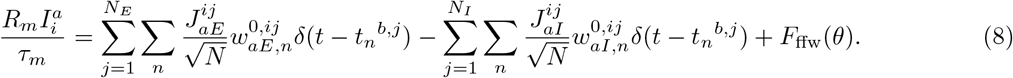

where Σ_*n*_ *δ*(*t* – *t_n_^b,j^*) is the spike train of the *j*-th neuron from population 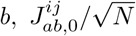 is the maximal synaptic weight from neuron *j* in population *b* to neuron *i* in population *a*, and 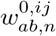. is a synaptic plasticity factor, which here depends on the spike time. If two neurons are connected, 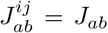, otherwise 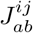 is zero. Given that each connection is re-scaled by 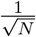, the total recurrent input is on the order of 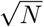. The feed-forward input is given by 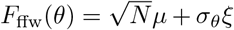. Here, 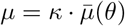, where *κ* ≥ 0 denotes contrast, 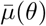 is the orientation-dependent component of the input, and *ξ* is white noise with standard deviation *σ_θ_*. All parameters are given in Table S1.

### Rate formalism

The firing rate of population *a* = {*E, I*} is given by *ν_a_*(*θ*) ≡ [〈*s_a,j_* (*t*)〉], where *s_a,j_*(*t*) ∑_*n*_ *δ*(*t* – *t_n_^a,j^*) is the spike train of the *j*-th neuron from population *a*, 〈·〉 denotes temporal average, and [·] denotes population average. Assuming E/I balance [46, 58, 36], the mean input currents are related to the firing rates as

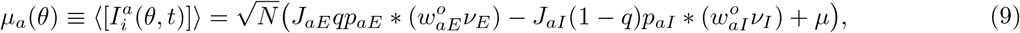

where * is the convolution in the orientation space, 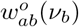 is a synaptic plasticity factor that modulates the effective synaptic strengths with the presynaptic activity, *p_aE_* and *p_aI_* are the connection probability functions between populations, and *μ* represents a stimulus of orientation *θ* and contrast *κ*. In the limit *N* → ∞ and requiring that *μ_α_* in Equation 9 is finite we get that

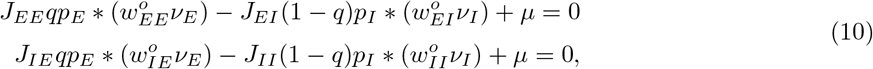

where we have made the assumption that *p_EE_* = *p_IE_* = *p_E_* and *p_EI_* = *p_II_* = *p_I_*. The probability of connection is given by 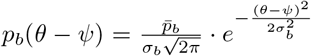 where *θ* – *ψ* denotes difference in preferred orientation and *σ_b_* denotes connectivity width. We model the input *μ* as a Gaussian function 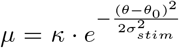, where *κ* is the stimulus contrast, *σ_stim_* is its tuning width, and *θ*_0_ is the stimulus orientation (see Table S1 for parameters).

### Network models

#### Model I

describes homogeneously randomly connected networks with constant synaptic weights (Figure S1). The connection probabilities *p_E_* and *p_I_* are uniform across the feature space Γ. The synaptic plasticity factors are constant and equal unity, 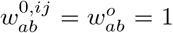. The external feed-forward input *F*_ffw_ is orientationindependent, with *μ* = *κ*.

#### Model II

describes homogeneously randomly connected networks with plastic *E* → *E* synaptic weights (Figure 3). The synaptic plasticity factor 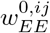 models STP as in Refs. [57, 36]. The firing rate approximation for 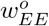 in STP is given in the Appendix. The synaptic plasticity factors for the remaining population connectivities are constant and equal unity. The connection probabilities *p_E_* and *p_I_* are constant. The external feed-forward input *F*_ffw_ is orientation-independent, with *μ* = *κ*.

#### Model III

describes networks of neurons connected with orientation-dependent probability and constant synaptic weights (Figure 4, Equation 4). For each neuron, *C_a_* presynaptic neurons are sampled from a probability distribution *p_ab_*(*θ* – *ψ*), which is described by a gaussian function of width 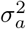. Therefore, the number of presynaptic inputs to a neuron for a particular network size *N* is the same as in Model I and II, but here the majority of connections are made within the nearby orientation space. The synaptic plasticity factors 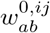 and 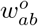 are constant and equal unity.

#### Model IV

describes networks of neurons connected with orientation-dependent probability and plastic *E* → *E* synaptic weights (Figure 5, Equation 5). The connectivity is implemented as in Model III, while the synaptic plasticity factors 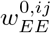 and 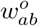 are defined as in Model II.

#### Model V

describes networks of neurons connected with orientation-dependent probability (see Model III) and with a power-law type of synaptic plasticity in the *E* → *E* and *E* → *I* connections (Figure 6, Equation 6):

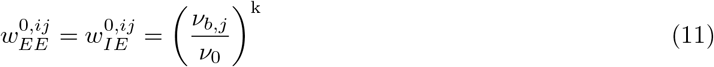

where *v*_0_ = 1 Hz, k denotes the synaptic efficacy. The firing rate *v_bj_* of neuron *j* is estimated using the sum of its last *n* =10 inter-spike intervals (ISIs) (see Appendix):

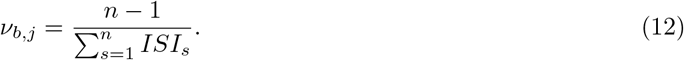

Parameters for each network model are given in Table S1.

### The susceptibility to input contrast *δ* in randomly connected networks with *E* → *E* plasticity

Here, we consider *E* → *E* plasticity (see Appendix for analogous calculations for networks with *E* → *E* and *E* → *I* plastic synapses). From Equation 10, we define the input contrast *κ* that is consistent with a balanced network with *E* → *E* plastic synapses to fire at rate 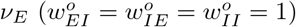

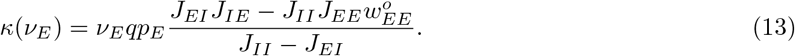

Using the definition of k_*EE*_ (Equation 3), the derivative of the input contrast *κ* with respect to the excitatory firing rate *v_E_* is inserted into Equation 1 to obtain

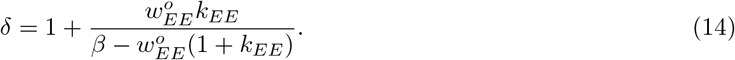

where 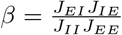. This equation is equivalent to Equation 2, where 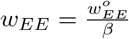. The phase space of values for *δ* is shown in Figure 2.

### Numerical methods and code availability

The spiking network code was written in C++ and the results analyzed in Matlab (Matlab 2018b, Mathworks). The code is available upon request.

## Acknowledgements

This work was funded by the Max Planck Society and the German Research Foundation via CRC 1080 (T.T.). A.N. and L.BT. acknowledge additional support from an add-on scholarship of the Joachim Herz Foundation. We thank Nataliya Kraynyukova, Carlos Wert Carvajal and Simon Renner for commenting on earlier versions of the manuscript, Alexander Dutine for validating the code, and members of the Tchumatchenko lab for fruitful discussions. S.K. acknowledges the support of the German Research Foundation (EXC 2181/1 - 390900948, the Heidelberg Excellence Cluster STRUCTURES). We acknowledge the support of the Center for Multiscale Modelling in Life Sciences (CMMS) by the LOEWE initiative of the Hessian government.

## Author contributions

L.BT. and T.T. conceived the study. L.BT. and S.K. developed the theoretical framework. L.BT. performed the analytic calculations, the numerical simulations and analyzed the data. P.E., A.N., and S.K. wrote the spiking network code. P.E., A.N., and L.BT. validated the code. L.BT. wrote the first draft of the manuscript. L.BT., A.N., P.E. and T.T wrote the manuscript. All authors provided critical feedback and helped shape the research and analysis.

## Competing interests

The authors declare no competing interests.

## Appendix

### Short-term plasticity model (STP)

Short-term plasticity is a non-linear synaptic mechanism observed in cortical pyramidal neurons [55, 57, 59, 22]. In these synapses, the synaptic strength, measured as the amplitude of the postsynaptic potential, depends on the availability of presynaptic vesicles and their release probability [57]. Tsodysks and Markram [57] first proposed a model of short-term plasticity, which was later modified by Mongillo et al. [36], who introduced synaptic state binary variables. Here, we briefly summarize this STP model in spiking networks. For more detailed information, we refer to Ref. [36]. We then provide details on the approximation we have used in this study to derive the mean-field steady-state synaptic factor *w*^o^.

Mongillo et al. [36] define STP variables *x* and *y*. At an individual synapse, *x* represents the availability of neurotransmitter and *y* is the binding state of calcium ions. Neurotransmitter can be available (*x* = 1) or not (*x* = 0) and calcium can be either bound (*y* = 1) or not (*y* = 0). If the presynaptic neuron spikes, calcium binds to the postsynaptic receptor with probability U. If calcium binds (*y* = 1) and neurotransmitter is available (*x* = 1), neurotransmitter is released (*x* → 1) and the postsynaptic neuron spikes. In between spikes, neurotransmitter replenishes with rate 1/*τ_D_* (*x* → 1) and calcium unbinds with rate 1/*τ_F_* (*y* → 0). In a network context, Mongillo et al. [36] define

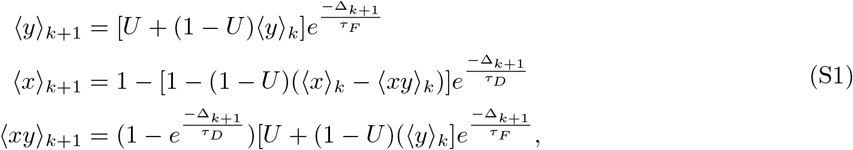

where 〈*y*〉 denotes the fraction of synapses with calcium being available, 〈*x*〉 the fraction of synapses with neurotransmitter being available, and 〈*xy*〉 the fraction of synapses where both calcium and neurotransmitter are available. The probability of neurotransmitter release upon the (*k* + 1)-th spike is *w*_*k*+1_ = **U**〈*x*〉_*k*+1_ + (1 – *U*)〈*xy*〉_*k*+1_. Mongillo et al. [36] compute the steady-state synaptic state as a function of the Laplace transform of the interspike interval probability distribution function. For arbitrary stationary interspike interval probability distribution functions, they derive the following steady-state values

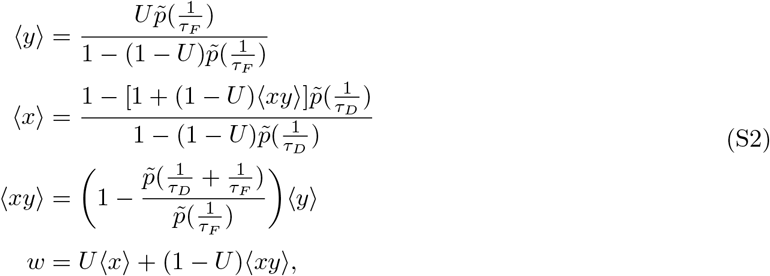

where 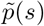 is the Laplace transform of the interspike interval probability distribution function and *w* is the steady-state neurotransmitter release probability upon spike.

At this point, we assume an exponential interspike interval distribution function, which is characteristic of a Poisson process [28], to get the steady state mean-field approximation for *w*. In this case, the Laplace transform is 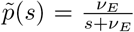, where *ν_E_* is the presynaptic excitatory firing rate. Inserting this expression into Equations (S2) gives the following approximations for the synaptic steady-states 〈*y*〉_*ν_E_*_, 〈*x*〉_*ν_E_*_, 〈*xy*〉_*ν_E_*_ and *w*^o^(*ν_E_*):

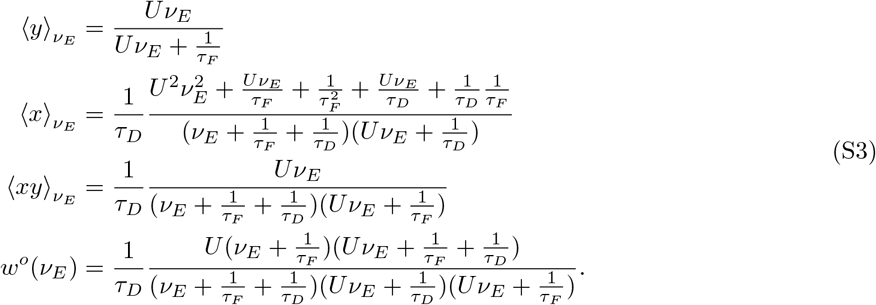

*w^o^*(*ν_E_*) is the steady-state probability of neurotransmitter release as a function of the presynaptic excitatory firing rate *ν_E_* and the synaptic parameters *τ_F_, τ_D_* and *U* (Figure S3). *w^o^*(*ν_E_*) is used in the mean-field description of networks with *E* → *E* STP (Model II, Model IV, Equation 5).

### The susceptibility *δ* in networks with *E* → *E* and *E* → *I* plastic synapses

Here, we expand the definition of *δ* in networks with *E* → *E* plasticity (Equation 2) to networks in which *E* → *E* and *E* → *I* synapses are plastic. We examine if the different types of contrast response described in Figure 2 for networks with *E* → *E* plasticity change through the introduction of *E* → *E* and *E* → *I* plastic synapses. To that end, we derive the stimulus contrast that drives a network to fire at rates *ν_E_* and *ν_I_* from Equation 10:

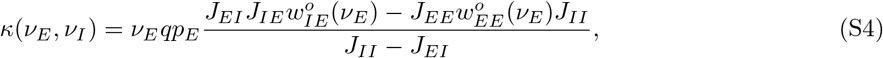

where we have set 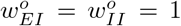 and k_*EI*_ = k_*II*_ = 0. Using the definition of k_*EE*_ (Equation 3), the derivative of the input contrast *κ* with respect to the excitatory firing rate *ν_E_* is inserted into Equation 1 to obtain

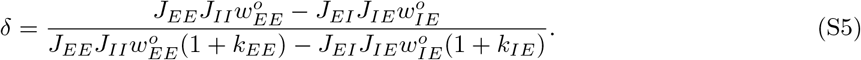

We substitute 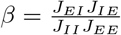, which yields

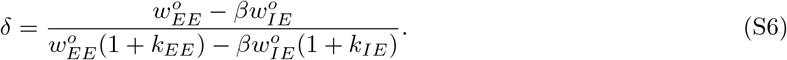

We examine the non-linear contrast response of balanced networks with additional plastic *E* → *I* synapses for *w_EE_* and *k_EE_* ≠ 0. The phase space of values of Equation S6 under these constraints is shown in Figure S2. Our results show that, similarly to the network with plastic *E* → *E* synapses (Figure 3), networks with both *E* → *E* and *E* → *I* plastic synapses can exhibit different types of non-linear contrast response. The type of nonlinearity (i.e sublinearity, supralinearity or supersaturation) is determined by the synaptic parameters 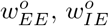, *k_EE_*, and *k_IE_*.

### Linear stability analysis

Our results show that plastic *E* → *E* synapses control the response to contrast in balanced networks *δ*. The contrast response function can be sublinear, supralinear or supersaturating depending on the synaptic parameters (Figures 2). Yet, the existence of these different steady states does not guarantee that they are stable in balanced networks. Here, we analyze the stability of the steady states in networks with plastic *E* → *E* synapses. Let us begin by assuming linear dynamics for *ν_E_*(*t*) and 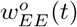 in the vicinity of the steady state. The excitatory firing rate *ν_E_* converges to the steady state 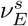 with timescale *τ_n_*. Similarly, the synaptic plasticity factor 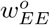 converges to the steady state value 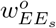 with timescale *τ_s_*. 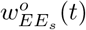 is determined by the network firing rate, such that 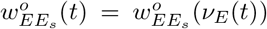. In cortical networks, the excitatory and inhibitory currents have been shown to correlate with milisecond precision [39]. In line with these observations, we assume that the firing rates balance instantaneously such that *τ_n_* ≪ *τ_s_*. This implies that (see Equation 13)

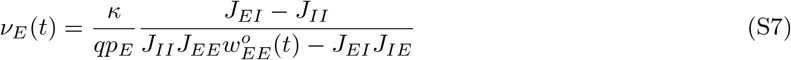

holds at all times *t*. With this, we are left with one differential equation for the synapses in the vicinity of the fixed points

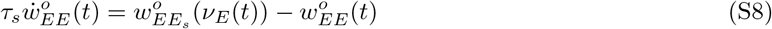

We compute 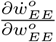 and evaluate it at the steady state 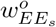. The sign of 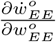 indicates the stability of the steady state. If 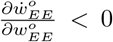, the steady state is stable as perturbations around the steady state are absorbed and the system is pushed back to the equilibrium point. If 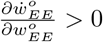, the steady state is unstable as perturbations around the steady state are amplified and the system is pushed away from the equilibrium point. From Equation S8, we obtain

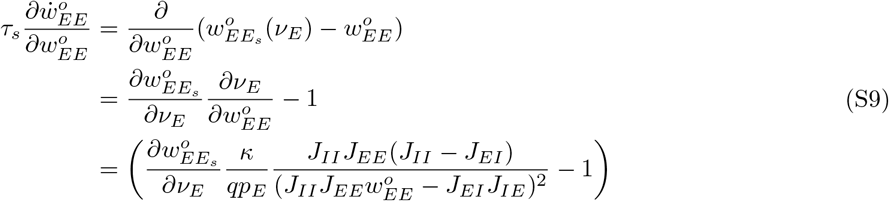

Next, we substitute *κ* by using the balanced state expression (see Equation S7). This yields

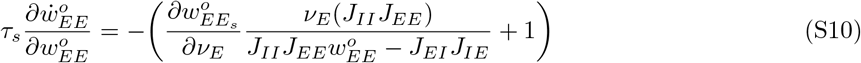

We now evaluate this expression for 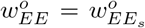 and make use of the definition of the synaptic efficacy k_*EE*_ introduced in the main text (Equation 3): here 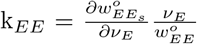. We substitute k_*EE*_ into Equation S10, which yields

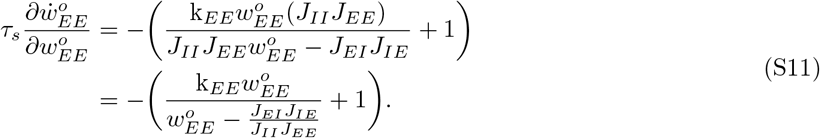

Using 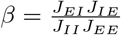, the above expression transforms into

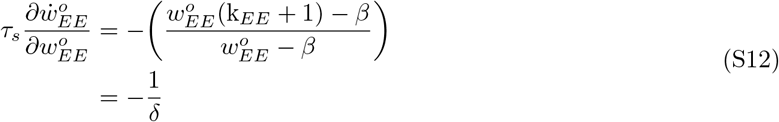

where *δ* is the susceptibility to input in balanced networks. From this, we conclude that the supersaturation regime (*δ* < 0) is not stable in the limit of balanced networks. Both sublinear and supralinear regimes (*δ* > 0) are stable.

### Estimation of the firing rate *ν* in Model V

In a spiking network simulation, the firing rate is usually obtained *a posteriori*, by dividing the number of spikes that occurred over a given time frame. In the context of power-law synapses, the firing rate needs to be estimated for each neuron, upon firing, in order to determine the amplitude of the post-synaptic potential:

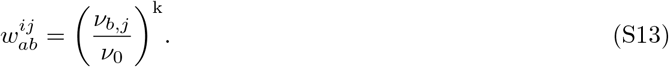

Typically, the firing rate of each neuron is estimated by counting the number of spikes emitted in a given period of time. This approach has multiple drawbacks. First, the number of spikes emitted in a given time period is discrete, so the relative precision of the firing rate estimation decreases for low firing rates. Second, the synapses would display limited responsiveness, as they cannot adapt to new states faster than the time needed to renew the time window. Finally, the choice of the optimal length of the time window is a trade-off between a better precision (long period of time) and responsiveness (short period of time). Making this trade-off is especially difficult in networks with feature-dependent connectivity and input, in which neurons of the same population have a wide range of firing rates. Instead, we choose to estimate the firing rates using the *n* interspike intervals (ISIs) that precede any new spike. Assuming that the neuron’s spiking follows a Poisson process and that the system is in steady state (all ISIs follow the same distribution), the sum of the *n* ISIs should follow an Erlang distribution

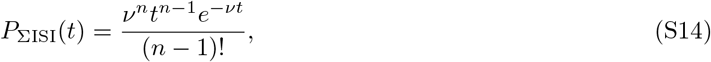

where ΣISI is the sum of *n* ISIs, and *ν* is the firing rate of the neurons. The mean of the n ISIs is given by

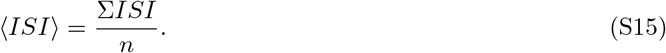

With the change of variable 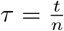

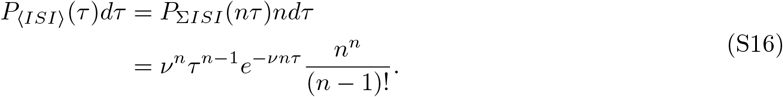

Following the same approach, the pdf of the inverse of the average ISI (〈*ISI*〉^−1^) can be obtained with the change of variable 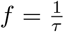

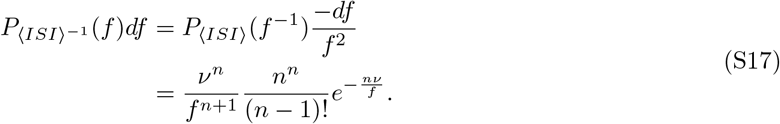

For *n* > 1, the expected value of 〈*ISI*〉^−1^ is given by

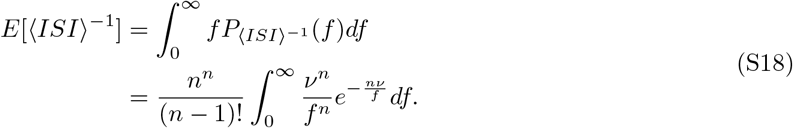

With one last change of variable 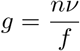

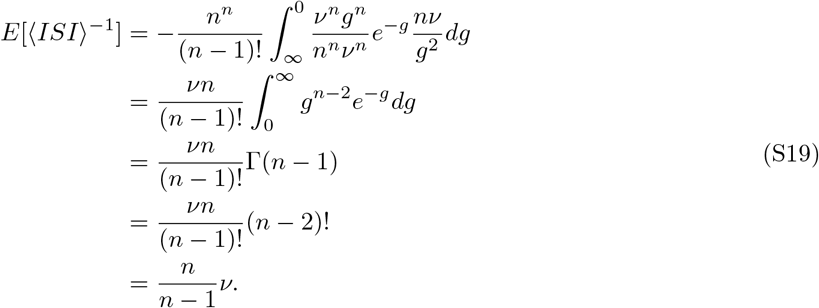

We can therefore build an estimator 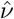 of the neuron’s firing rate based on the *n* ISIs that precede the time of evaluation:

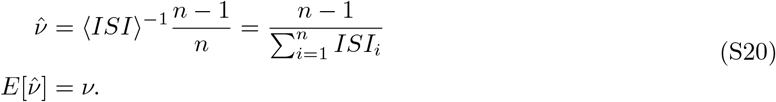

### Firing rate profile *ν_E_* of balanced networks with feature-dependent connectivity

In this section, we outline a rate formalism of balanced networks with feature-dependent connectivity and plastic synaptic weights.

The firing rate of the presynaptic population *a* = {*E, I*} is given by *ν_α_*(*θ*) ≡ [〈*s_a,i_*(*t*)〉], where 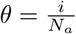 and *s_a,i_*(*t*) = *δ*(*t* – *t_n_^a,i^*) is the spike train of the *i*-th neuron from population *a*, and *t_n_^a,j^* are its spike times. In the continuum limit, the mean input currents are related to the firing rates as follows

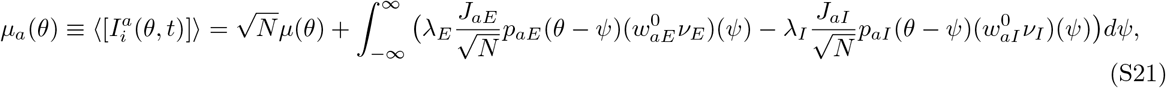

where 〈·〉 denotes temporal average, [·] denotes population average. The synaptic plasticity factor 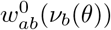 modulates the synaptic strength *J_ab_* as a function of the presynaptic activity. The term *μ*(*θ*) denotes a feature-dependent external input, λ_*b*_ is the linear density of population b in the feature space. Since we assume the feature space Γ to have length 1, we have λ_*b*_ = *N_b_*.

For *μ_a_*(*θ*) to be finite, the following condition must be met:

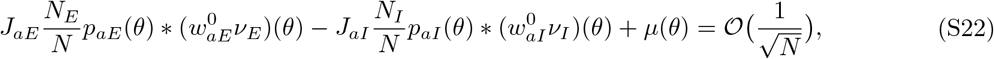

where * is the convolution in the feature space. Taking the limit *N* → ∞ and writing the resulting equation in the Fourier domain we get

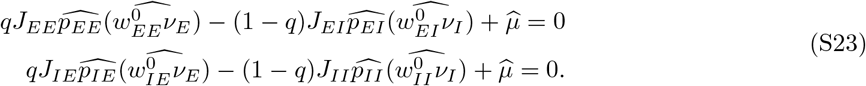

Assuming the inhibitory connections are constant 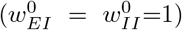, and setting *p_EE_* = *p_IE_* = *p_E_* and *p_EI_* = *p_II_* = *p_I_* the balanced state solution in the Fourier domain yields

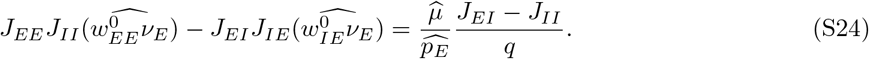

We set the feedforward input *μ*(*θ*) to be a gaussian function of preferred feature:

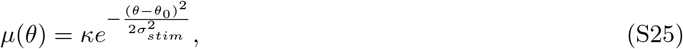

where *κ* is the stimulus contrast, 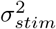 is the tuning width of the feedforward input and *θ*_0_ is the input feature.

Neurons with a similar preferred feature are more likely to be connected. We assume it is given as a Gaussian function:

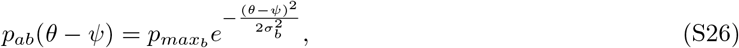

where *p_max_b__* is the peak probability of connection, for two neurons sharing the same feature preference, *θ* and *ψ* are the feature preference of the postsynaptic and presynaptic neurons, and *σ_b_* is the width of the distribution. The number of connections *C_b_* from population b is given by

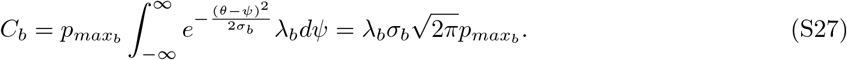

From this, we can define the connection probability with respect to the presynaptic population, such that *C_b_* = *p_b_N_b_*:

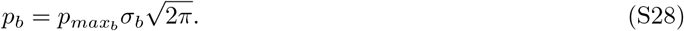

The excitatory firing rate profile can be expressed in the Fourier domain as

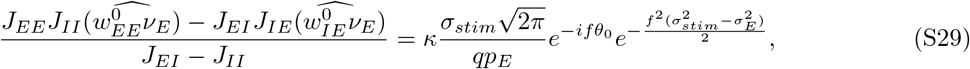

and can be transformed back into the feature space:

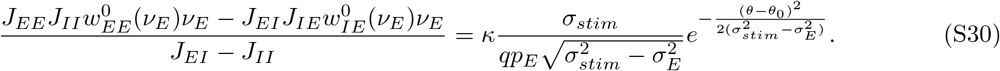

If the synaptic plasticity factor in *E* → *E* connections is the same as in *E* → *I* connections, the previous expression can be simplified as

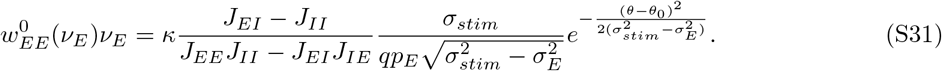

Finally, if the synaptic plasticity follows a power law, 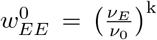, the firing rate response can be factorized into a function of contrast and a function of orientation:

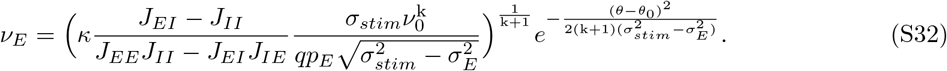

## Supplementary tables and figures

**Table S1:** Parameters for each network model. For all spiking network models: *τ_m_* = 20 (ms), *V_th_* = 1 (mV), *v_rest_* = 0 (mV), 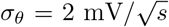, the simulation time step dt = 0.05 (ms), and the recording bin size is 50 ms. The synaptic weights and the input are re-scaled by 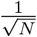 and 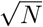, respectively (see Equation 8) [58]. Let us note that we re-scale the feature space Γ, *σ_E_* and *σ_I_* to 180° in the figures for illustration purposes.

| Model I |  |
| --- | --- |
| Populations | $N_E = [1.6 \times 10^4 \dots 6 \times 10^4]$ ; $N_I = [4 \times 10^4 \dots 1.5 \times 10^4]$ (neurons) |
| Connectivity | Random, $p_E = \bar{p}_E = 0.05$; $p_I = \bar{p}_I = 0.05$ |
| Synaptic weights | $J_{EE} = 2.5$; $J_{EI} = 10$; $J_{IE} = 4$; $J_{II} = 13.5$ (mV/spike) |
| Synaptic plasticity | No |
| Input | Orientation-independent, $\kappa = [0 \dots 2.5]$ (mV/s) |
| Model II |  |
| Populations | $N_E = 10^5$; $N_I = 2.5 \times 10^4$ (neurons) |
| Connectivity | Random, $p_E = \bar{p}_E = 0.05$, $p_I = \bar{p}_I = 0.05$ |
| Synaptic weights | $J_{EE} = 8$; $J_{EI} = 10$; $J_{IE} = 4$; $J_{II} = 13.5$ (mV/spike) |
| Synaptic plasticity | STP in $E \rightarrow E$:<br>Facilitation: $U = 0.05$, $\tau_F = 0.8$ ms, $\tau_D = 0.03$ ms<br>Depression: $U = 0.35$, $\tau_F = 0.15$ ms, $\tau_D = 0.7$ ms |
| Input | Orientation-independent, $\kappa = [0 \dots 2.5]$ (mV/s) |
| Model III |  |
| Populations | $N_E = 2 \times 10^5$; $N_I = 5 \times 10^4$ (neurons) |
| Connectivity | Orientation-dependent, $\bar{p}_E = \bar{p}_I = 0.05$; $\sigma_E = \sigma_I = 0.1$ |
| Synaptic weights | $J_{EE} = 2.5$; $J_{EI} = 10$; $J_{IE} = 4$; $J_{II} = 13.5$ (mV/spike) |
| Synaptic plasticity | No |
| Input | Tuned gaussian input, $\kappa = 1, 2, 3$ (mV/s), $\sigma_{stim} = 0.16$ |
| Model IV |  |
| Populations | $N_E = 4 \times 10^5$; $N_I = 10^5$ (neurons) |
| Connectivity | Orientation-dependent, $\bar{p}_E = \bar{p}_I = 0.05$; $\sigma_E = \sigma_I = 0.1$ |
| Synaptic weights | $J_{EE} = 8$; $J_{EI} = 10$; $J_{IE} = 4$; $J_{II} = 13.5$ (mV/spike) |
| Synaptic plasticity | STP in $E \rightarrow E$:<br>Facilitation: $U = 0.05$, $\tau_F = 0.8$ ms, $\tau_D = 0.03$ ms<br>Depression: $U = 0.2$, $\tau_F = 0.2$ ms, $\tau_D = 0.2$ ms |
| Input | Tuned gaussian input, $\kappa = [0 \dots 4]$ (mV/s), $\sigma_{stim} = 0.16$ |
| Model V |  |
| Populations | $N_E = 4 \times 10^5$; $N_I = 10^5$ (neurons) |
| Connectivity | Orientation-dependent, $\bar{p}_E = \bar{p}_I = 0.05$; $\sigma_E = \sigma_I = 0.1$ |
| Synaptic weights | $J_{EE} = 2$; $J_{EI} = 10$; $J_{IE} = 5$; $J_{II} = 12$ (mV/spike) |
| Synaptic plasticity | Power-law plasticity in $E \rightarrow E$ and $E \rightarrow I$:<br>Synaptic efficacy $k = -0.5$ |
| Input | Tuned gaussian input, $\kappa = 0.5, 1, 1.5, 2$ (mV/s), $\sigma_{stim} = 0.16$ |

**Figure S1:**
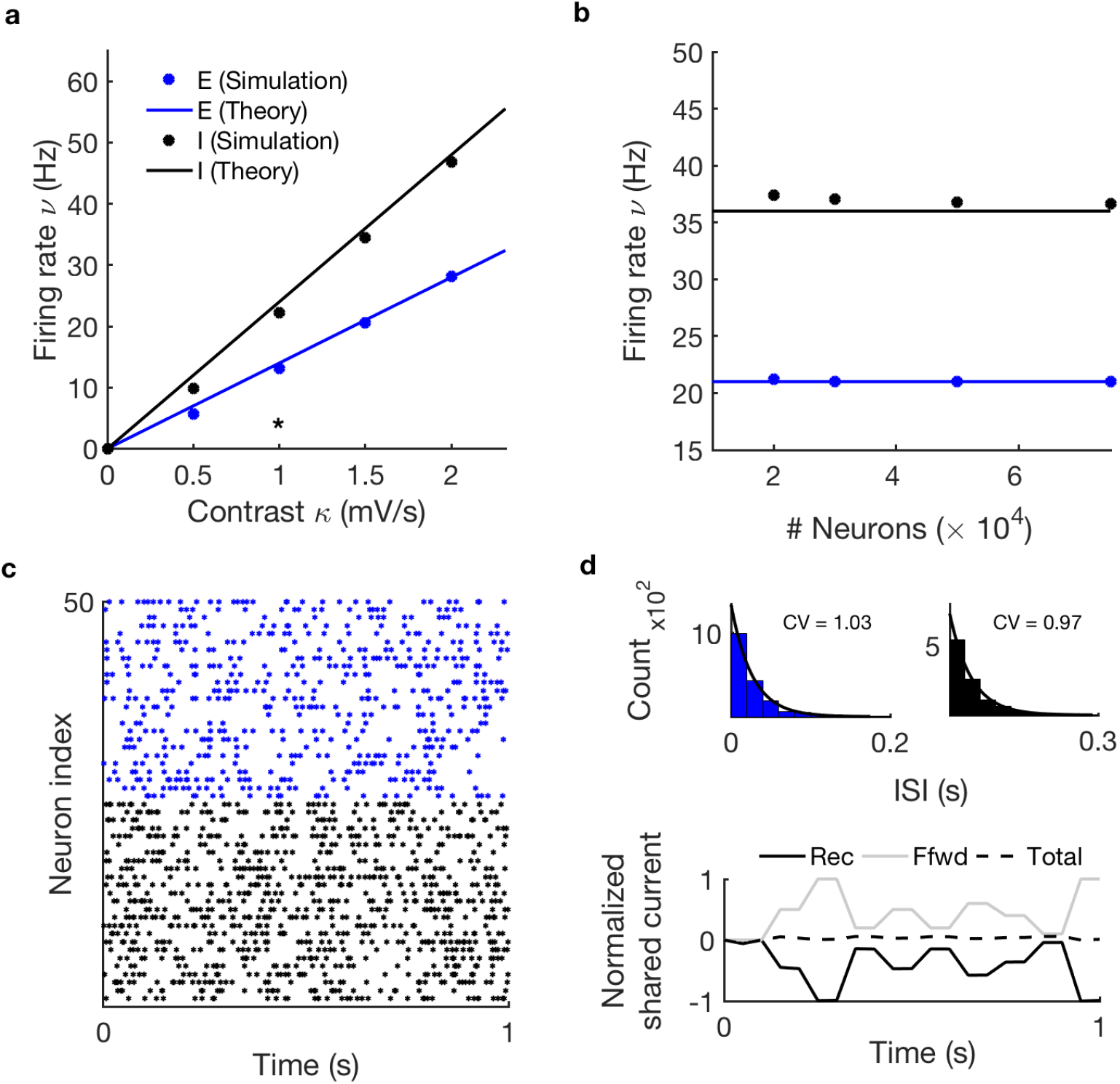
Linear response to contrast for homogeneous networks of randomly connected LIF neurons and constant synapses. **(a)** The mean activity of the excitatory (blue) and the inhibitory (black) populations in a network of *N* = 5 · 10^4^ LIF neurons (dots) with constant synapses 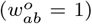 is captured by the balanced state equations (Equation 10, solid lines). **(b)** An increase in the number of neurons from *N* = 2 · 10^4^ to *N* = 5 · 10^4^ corrects a maximum deviation from balance (solid lines) of 4.5%. **(c)** Spike raster of 25 excitatory and 25 inhibitory neurons across 1 s for the data point denoted by an asterisk in (a). **(d)** Top: coefficient of variation (CV), inter-spike interval (ISI) distribution and exponential fit for the data point denoted by asterisk in (a) indicate asynchronous irregular firing. Bottom: temporal correlations of the E and I inputs estimated for data point denoted by asterisk in (a) indicate E-I balance. For parameters see Table S1, Model I.

**Figure S2:**
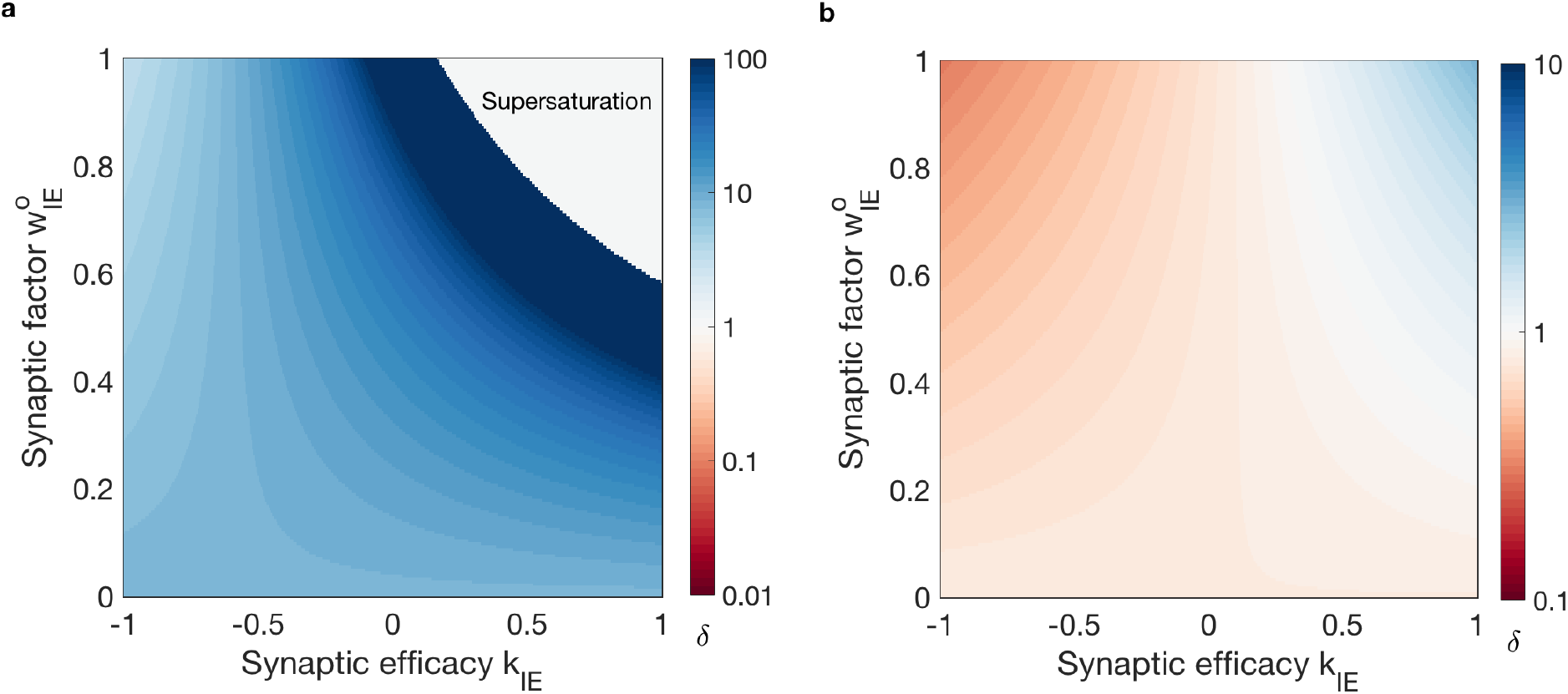
Susceptibility to contrast *δ* in networks with plastic *E* → *E* and *E* → *I* synapses. **(a)** The phase space of values for *δ* (Equation S6) for 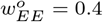 and k_*EE*_ = −0.6 (depressing state). **(b)** The phase space of values for *δ* (Equation S6) for 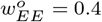 and k_*EE*_ = 0.1 (facilitating state). Let us note that for positive input contrast and *J_II_* > *J_EI_*, parameters must satisfy 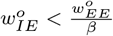 for positive firing rates [58, 46].

**Figure S3:**
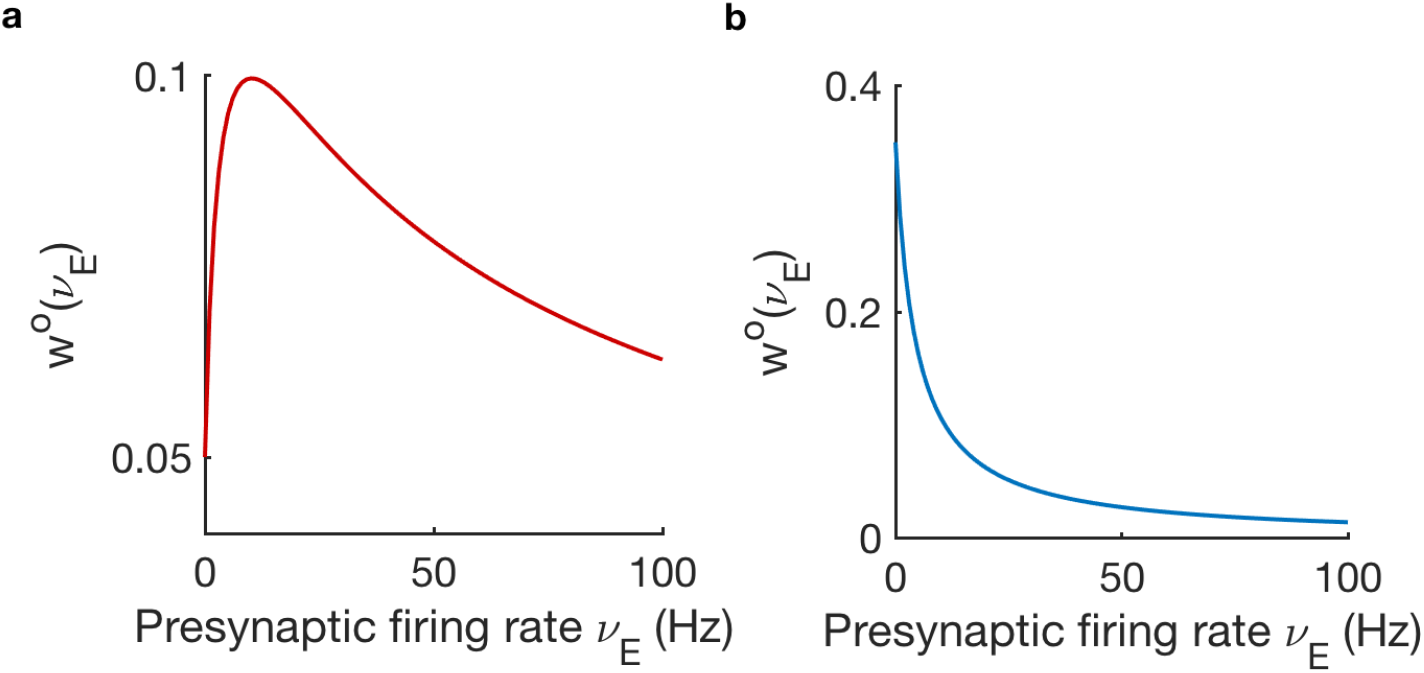
Steady-state probability of neurotransmitter release *w^o^*(*ν_E_*) (Equation S3). **(a)** Facilitating transmission: *w^o^* increases until neurotransmitter vesicles can not replenish fast enough to be released upon spike arrival. At this point, *w^o^* decreases for increasing *ν_E_*. Parameters: *U* = 0.05, *τ_F_* = 0.8 ms, *τ_D_* = 0.3 ms. This function models the *E* → *E* synapses in Figures 2,3 and 5. **(b)** Depressing transmission: *w^o^* decreases with increasing *ν_E_* as a result of neurotransmitter not being available. Parameters: *U* = 0.35, *τ_F_* = 0.15 ms, *τ_D_* = 0.7 ms. This function models the *E* → *E* synapses in Figures 2 and 3.

**Figure S4:**
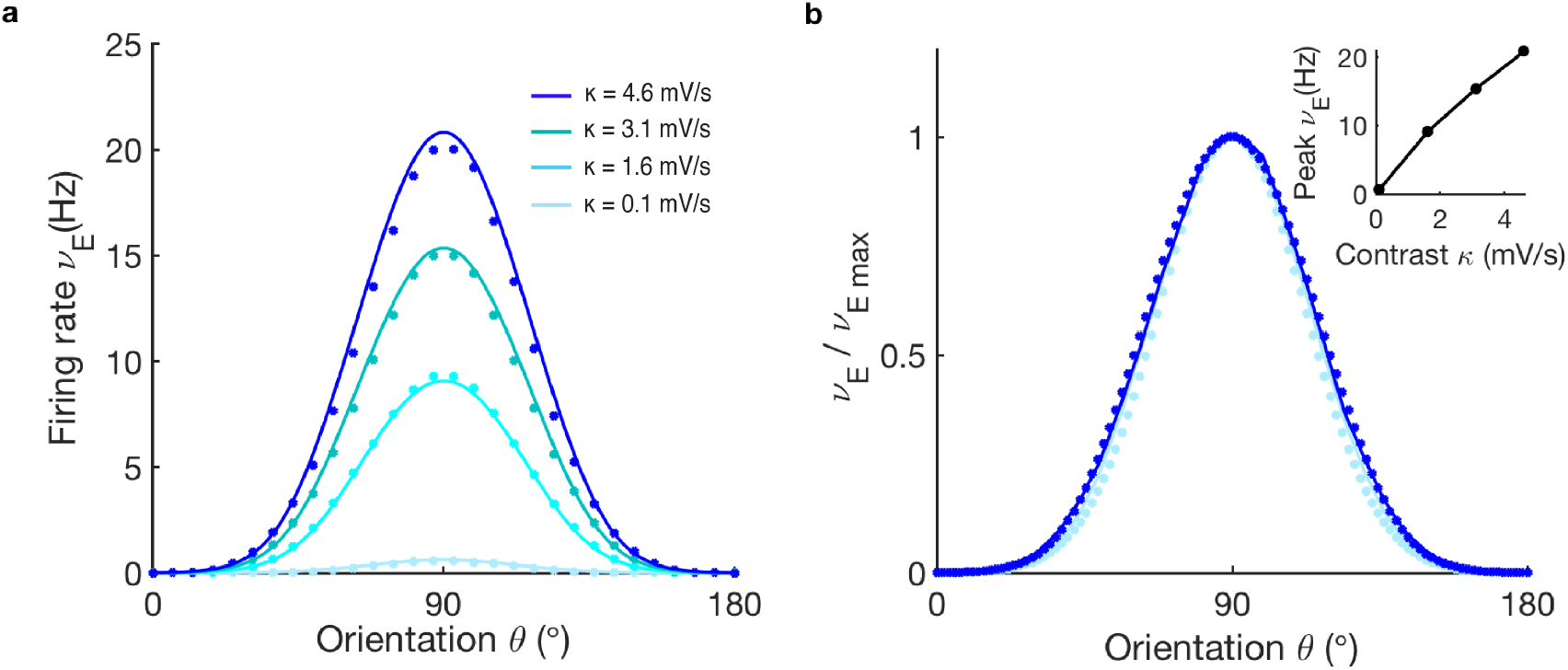
Depressing STP in the *E* → *E* synapses induces sublinear susceptibility to input contrast and slightly broadens the network selectivity in balanced networks. Same network model as in Figure 5a with depressing *E* → *E* STP synapses (see Methods - Model IV and Table S1). **(a)** Excitatory tuning curves for different input contrast, *κ* = 0.1,1.6, 3.1,4.6 (mV/s), in a mean-field network description (Equation 5, solid lines) compared to spiking network simulations of N = 5 · 10^5^ neurons (dots). Notice the difference in tuning curves compared to a network with facilitating STP *E* → *E* synapses (Figure 5). **(b)** Normalized excitatory tuning curves in (a) show a slight decrease in selectivity as a function of input contrast. Inset: firing rate of tuning curves in (a) in spiking networks (dots) and balanced theory (solid line) for *θ* = *θ*_0_. Synaptic parameters: *U* = 0.2, *τ_F_* = 0.2 ms, *τ_D_* = 0.2 ms.

**Figure S5:**
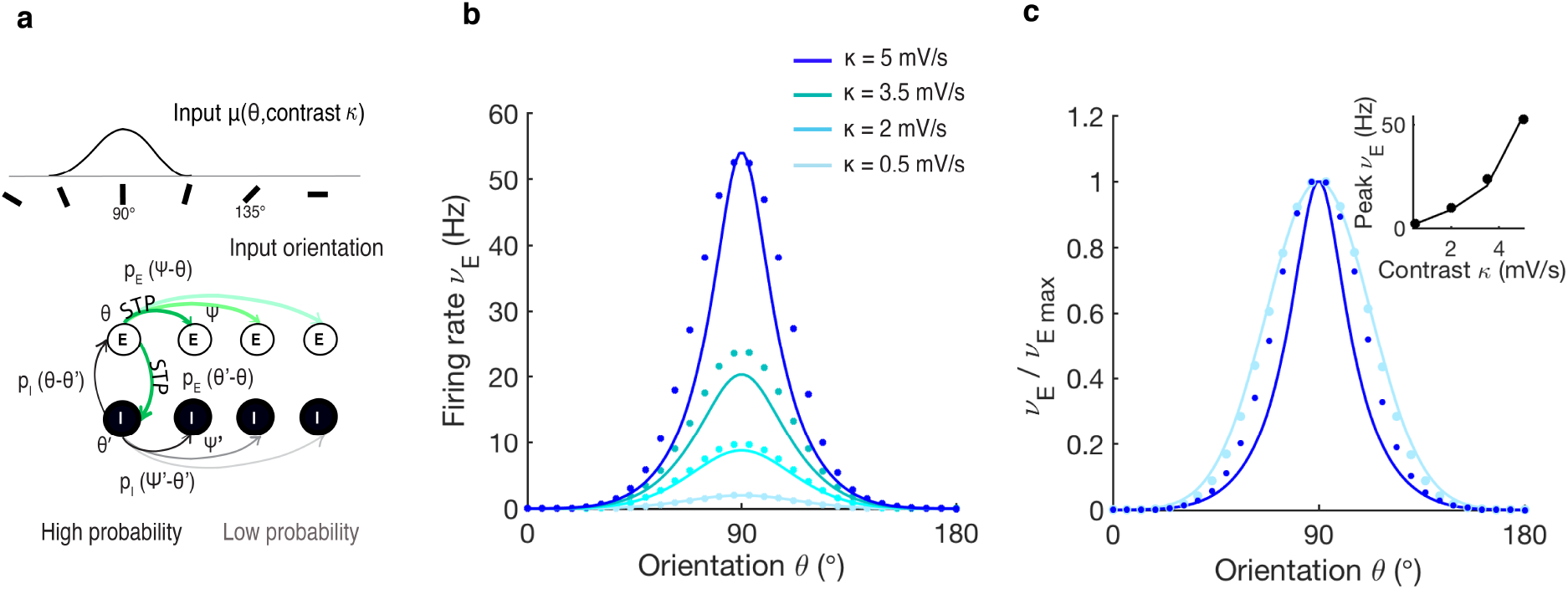
Depressing STP in the *E* → *E* and *E* → *I* synapses induces a supralinear response to contrast and narrows the network selectivity. **(a)** Network with orientation-dependent connectivity and *E* → *E* and *E* → *I* STP synapses (parameters as in Model IV, see Table S1). **(b)** Excitatory tuning curves for different input contrast, *κ* = 0.5, 2, 3.5, 5 (mV/s), in a mean-field network description (Equation 10, solid lines) compared to spiking network simulations of *N* = 5 · 10^5^ neurons (dots). **(c)** Normalized excitatory tuning curves for *κ* = 0.5 and *κ* = 5 (mV/s) show the increase in selectivity as a function of input contrast, color code as in (b). Inset: peak firing rate of tuning curves in (b) in spiking networks (dots) and balanced theory (solid line) for *θ* = *θ*_0_. Synaptic weights *J_EE_* = 10; *J_EI_* = 10; *J_IE_* = 27; *J_II_* = 13.5 (mV/spike). Depression parameters for *E* → *E* and *E* → *I* synapses: *U* = 0.2, *τ_F_* = 0.2 and *τ_D_* = 0.2.

**Figure S6:**
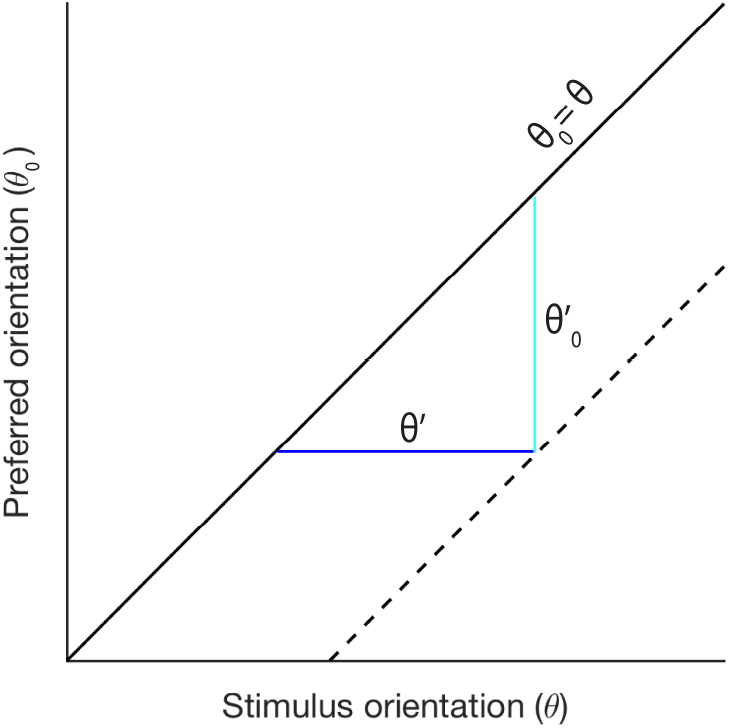
The network response depends on the difference between the neuron’s preference and the stimulus orientation. Previous approaches have defined the network response to orientation *ν* as the mean activity of single neurons aligned at the preferred orientation [8]. This can be mathematically formalized as 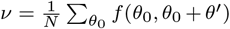. We have assumed a preferred orientation for each neuron and defined the network response as the mean activity of the neurons relative to the stimulus orientation. Mathematically, this implies that 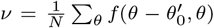. Here we show that both descriptions of the network response are equivalent as *θ* = *θ*_0_ + *θ*′.

**Video S1:**
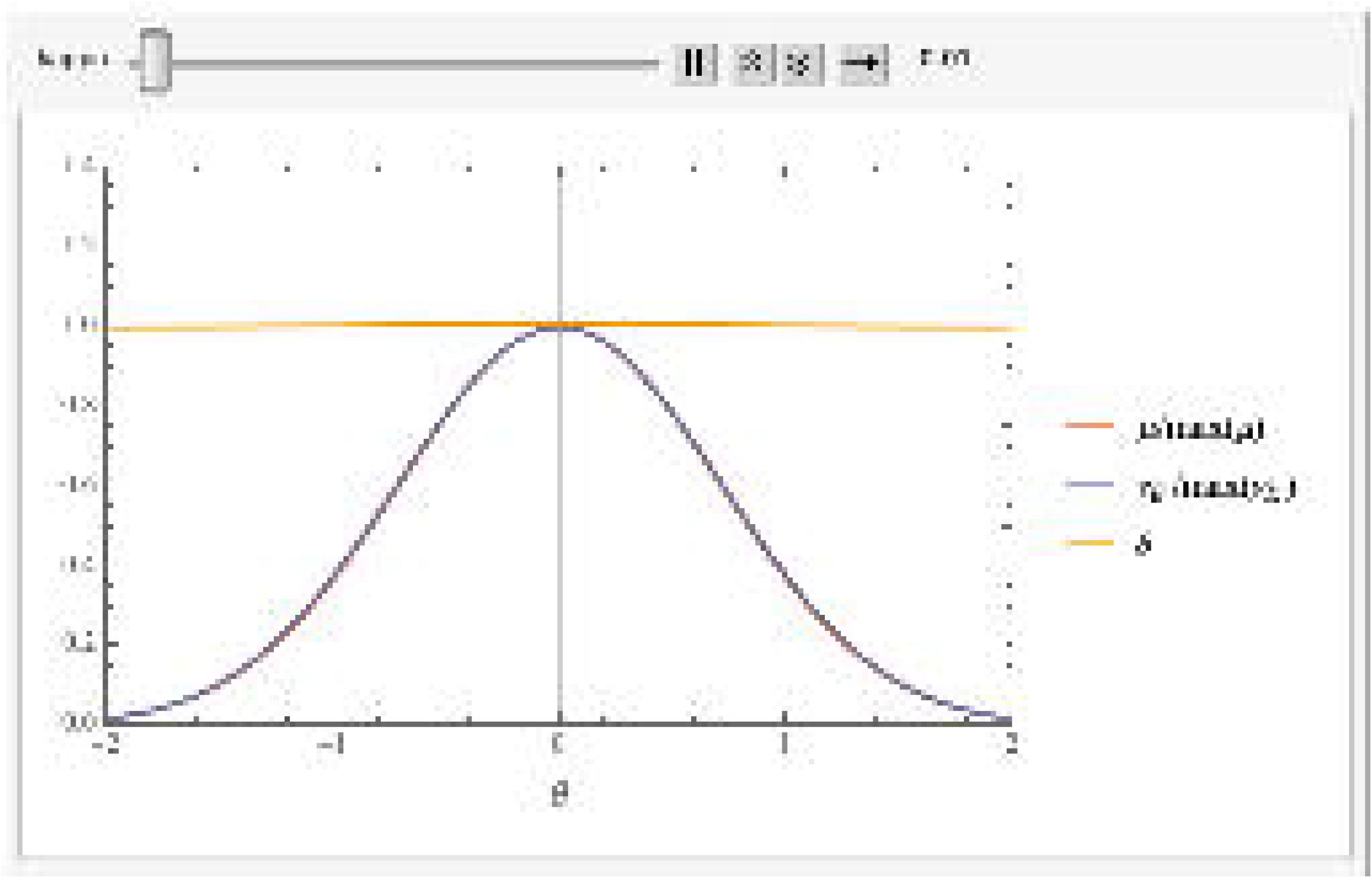
Approximation for excitatory firing rate *ν_E_* in a network with *E* → *E* plasticity with feature-dependent input *μ*. Here, *ν_E_* = *μ*^*δ*(*μ*)^ is used, where for simplicity dimensionless quantities are used, and where we have used *δ*(*μ*) as a first order approximation for *δ*(*ν_E_*). This is a valid approximation for *δ* close to unity. Here, *μ* = *κe*^−*θ*^2^^ and preferred orientation *θ* = 0. As a plasticity rule, an STP-like rule *w*(*ν*) = (1 – *e*^−*ν*^)*e*^−0.3*ν*^ is used, where the term (1 – *e*^−*ν*^) implements facilitation and the subsequent term *e*^−0.3*ν*^ implements depression. Equation 1 is then used to derive the expression for *δ*(*ν*). One can observe a regime for small contrasts *κ* < 1 in which *δ* > 1 in the preferred orientation, and in which *ν_E_* narrows. This is followed by a regime *κ* > 1 in which *δ* in the preferred orientation drops below unity, and the output firing rate broadens.

## Notes

### Competing Interest Statement

The authors have declared no competing interest.

## References

[1] Larry F Abbott, JA Varela, Kamal Sen, and SB Nelson. Synaptic depression and cortical gain control. Science, 275(5297):221–224, 1997.

[2] Yashar Ahmadian, Daniel B Rubin, and Kenneth D Miller. Analysis of the stabilized supralinear network. Neural computation, 25(8):1994–2037, 2013.

[3] Duane G Albrecht and David B Hamilton. Striate cortex of monkey and cat: contrast response function. Journal of neurophysiology, 48(1):217–237, 1982.

[4] Jeffrey S Anderson, Matteo Carandini, and David Ferster. Orientation tuning of input conductance, excitation, and inhibition in cat primary visual cortex. Journal of neurophysiology, 84(2):909–926, 2000.

[5] Kevin A Bolding and Kevin M Franks. Recurrent cortical circuits implement concentration-invariant odor coding. Science, 361(6407):eaat6904, 2018.

[6] J Gerard G Borst. The low synaptic release probability in vivo. Trends in neurosciences, 33(6):259–266, 2010.

[7] Tiago Branco and Kevin Staras. The probability of neurotransmitter release: variability and feedback control at single synapses. Nature Reviews Neuroscience, 10(5):373–383, 2009.

[8] Laura Busse, Alex R Wade, and Matteo Carandini. Representation of concurrent stimuli by population activity in visual cortex. Neuron, 64(6):931–942, 2009.

[9] Matteo Carandini. Amplification of trial-to-trial response variability by neurons in visual cortex. PLoS biology, 2(9):e264, 2004.

[10] Matteo Carandini and Frank Sengpiel. Contrast invariance of functional maps in cat primary visual cortex. Journal of vision, 4(3):1–1, 2004.

[11] Li Chao-Yi and O Creutzfeldt. The representation of contrast and other stimulus parameters by single neurons in area 17 of the cat. Pflügers Archiv, 401(3):304–314, 1984.

[12] H Cheng, Yuzo M Chino, EL Smith 3rd, Junji Hamamoto, and Kazuyuki Yoshida. Transfer characteristics of lateral geniculate nucleus x neurons in the cat: effects of spatial frequency and contrast. Journal of neurophysiology, 74(6):2548–2557, 1995.

[13] Sooyoung Chung and David Ferster. Strength and orientation tuning of the thalamic input to simple cells revealed by electrically evoked cortical suppression. Neuron, 20(6):1177–1189, 1998.

[14] Rodney J Douglas and Kevan AC Martin. Neuronal circuits of the neocortex. Annu. Rev. Neurosci., 27:419–451, 2004.

[15] David Ferster, Sooyoung Chung, and Heidi Wheat. Orientation selectivity of thalamic input to simple cells of cat visual cortex. Nature, 380(6571):249–252, 1996.

[16] David Ferster and Kenneth D Miller. Neural mechanisms of orientation selectivity in the visual cortex. Annual review of neuroscience, 23(1):441–471, 2000.

[17] Ian M Finn, Nicholas J Priebe, and David Ferster. The emergence of contrast-invariant orientation tuning in simple cells of cat visual cortex. Neuron, 54(1):137–152, 2007.

[18] Soledad Gonzalo Cogno and Germán Mato. The effect of synaptic plasticity on orientation selectivity in a balanced model of primary visual cortex. Frontiers in neural circuits, 9:42, 2015.

[19] D Hansel and C Van Vreeswijk. How noise contributes to contrast invariance of orientation tuning in cat visual cortex. Journal of Neuroscience, 22(12):5118–5128, 2002.

[20] David Hansel and German Mato. Short-term plasticity explains irregular persistent activity in working memory tasks. Journal of Neuroscience, 33(1): 133–149, 2013.

[21] David Hansel and Carl van Vreeswijk. The mechanism of orientation selectivity in primary visual cortex without a functional map. Journal of Neuroscience, 32(12): 4049–4064, 2012.

[22] Chris M Hempel, Kenichi H Hartman, X-J Wang, Gina G Turrigiano, and Sacha B Nelson. Multiple forms of short-term plasticity at excitatory synapses in rat medial prefrontal cortex. Journal of neurophysiology, 83(5): 3031–3041, 2000.

[23] David H Hubel and Torsten N Wiesel. Receptive fields, binocular interaction and functional architecture in the cat’s visual cortex. The Journal of physiology, 160(1): 106–154, 1962.

[24] Jean-Sébastien Jouhanneau, Jens Kremkow, Anja L Dorrn, and James FA Poulet. In vivo monosynaptic excitatory transmission between layer 2 cortical pyramidal neurons. Cell reports, 13(10): 2098–2106, 2015.

[25] Jean-Séebastien Jouhanneau and James FA Poulet. Multiple two-photon targeted whole-cell patch-clamp recordings from monosynaptically connected neurons in vivo. Frontiers in synaptic neuroscience, 11:15, 2019.

[26] Michael I Kerr, Mark J Wall, and Magnus JE Richardson. Adenosine a1 receptor activation mediates the developmental shift at layer 5 pyramidal cell synapses and is a determinant of mature synaptic strength. The Journal of physiology, 591(13): 3371–3380, 2013.

[27] Ho Ko, Sonja B Hofer, Bruno Pichler, Katherine A Buchanan, P Jesper Sjöström, and Thomas D Mrsic-Flogel. Functional specificity of local synaptic connections in neocortical networks. Nature, 473(7345): 87–91, 2011.

[28] Lubomir Kostal, Petr Lansky, and Michael Stiber. Statistics of inverse interspike intervals: The instantaneous firing rate revisited. Chaos: An Interdisciplinary Journal of Nonlinear Science, 28(10):106305, 2018.

[29] Robert B Levy and Alex D Reyes. Spatial profile of excitatory and inhibitory synaptic connectivity in mouse primary auditory cortex. Journal of Neuroscience, 32(16): 5609–5619, 2012.

[30] Jennifer F Linden, Robert C Liu, Maneesh Sahani, Christoph E Schreiner, and Michael M Merzenich. Spectrotemporal structure of receptive fields in areas ai and aaf of mouse auditory cortex. Journal of neurophysiology, 90(4):2660–2675, 2003.

[31] Eve Marder. Neuromodulation of neuronal circuits: back to the future. Neuron, 76(1):1–11, 2012.

[32] Thomas M McKenna, John H Ashe, and Norman M Weinberger. Cholinergic modulation of frequency receptive fields in auditory cortex: I. frequency-specific effects of muscarinic agonists. Synapse, 4(1):30—43, 1989.

[33] Douglas L Meinecke and Alan Peters. Gaba immunoreactive neurons in rat visual cortex. Journal of Comparative Neurology, 261(3):388–404, 1987.

[34] Raju Metherate, Nicole Tremblay, and Robert W Dykes. The effects of acetylcholine on response properties of cat somatosensory cortical neurons. Journal of Neurophysiology, 59(4):1231–1252, 1988.

[35] Kenneth D Miller and Todd W Troyer. Neural noise can explain expansive, power-law nonlinearities in neural response functions. Journal of neurophysiology, 87(2):653–659, 2002.

[36] Gianluigi Mongillo, David Hansel, and Carl Van Vreeswijk. Bistability and spatiotemporal irregularity in neuronal networks with nonlinear synaptic transmission. Physical review letters, 108(15):158101, 2012.

[37] Dina Moshitch, Liora Las, Nachum Ulanovsky, Omer Bar-Yosef, and Israel Nelken. Responses of neurons in primary auditory cortex (a1) to pure tones in the halothane-anesthetized cat. Journal of neurophysiology, 95(6):3756–3769, 2006.

[38] Ian Nauhaus, Laura Busse, Matteo Carandini, and Dario L Ringach. Stimulus contrast modulates functional connectivity in visual cortex. Nature neuroscience, 12(1):70, 2009.

[39] Michael Okun and Ilan Lampl. Instantaneous correlation of excitation and inhibition during ongoing and sensory-evoked activities. Nature neuroscience, 11(5):535, 2008.

[40] Aurélie Pala and Carl CH Petersen. In vivo measurement of cell-type-specific synaptic connectivity and synaptic transmission in layer 2/3 mouse barrel cortex. Neuron, 85(1): 68–75, 2015.

[41] Nicholas J Priebe and David Ferster. Inhibition, spike threshold, and stimulus selectivity in primary visual cortex. Neuron, 57(4): 482–497, 2008.

[42] Nicholas J Priebe, Ferenc Mechler, Matteo Carandini, and David Ferster. The contribution of spike threshold to the dichotomy of cortical simple and complex cells. Nature neuroscience, 7(10):1113, 2004.

[43] Guanxiao Qi, Karlijn van Aerde, Ted Abel, and Dirk Feldmeyer. Adenosine differentially modulates synaptic transmission of excitatory and inhibitory microcircuits in layer 4 of rat barrel cortex. Cerebral Cortex, 27(9):4411–4422, 2017.

[44] Gregg H Recanzone, Darren C Guard, and Mimi L Phan. Frequency and intensity response properties of single neurons in the auditory cortex of the behaving macaque monkey. Journal of neurophysiology, 83(4):2315–2331, 2000.

[45] AJ Rockel, Robert W Hiorns, and TP Powell. The basic uniformity in structure of the neocortex. Brain: a journal of neurology, 103(2): 221–244, 1980.

[46] Robert Rosenbaum and Brent Doiron. Balanced networks of spiking neurons with spatially dependent recurrent connections. Physical Review X, 4(2): 021039, 2014.

[47] Daniel B Rubin, Stephen D Van Hooser, and Kenneth D Miller. The stabilized supralinear network: a unifying circuit motif underlying multi-input integration in sensory cortex. Neuron, 85(2):402–417, 2015.

[48] Sadra Sadeh, Claudia Clopath, and Stefan Rotter. Emergence of functional specificity in balanced networks with synaptic plasticity. PLoS computational biology, 11(6): e1004307, 2015.

[49] Alessandro Sanzeni, Mark H Histed, and Nicolas Brunel. Response nonlinearities in networks of spiking neurons. PLoS computational biology, 16(9): e1008165, 2020.

[50] Christoph E Schreiner. Spatial distribution of responses to simple and complex sounds in the primary auditory cortex. Audiology and Neurotology, 3(2-3):104–122, 1998.

[51] Christoph E Schreiner, Heather L Read, and Mitchell L Sutter. Modular organization of frequency integration in primary auditory cortex. Annual review of neuroscience, 23(1): 501–529, 2000.

[52] G Sclar and RD Freeman. Orientation selectivity in the cat’s striate cortex is invariant with stimulus contrast. Experimental brain research, 46(3): 457–461, 1982.

[53] William R Softky and Christof Koch. The highly irregular firing of cortical cells is inconsistent with temporal integration of random epsps. Journal of Neuroscience, 13(1): 334–350, 1993.

[54] Andrew YY Tan, Li I Zhang, Michael M Merzenich, and Christoph E Schreiner. Tone-evoked excitatory and inhibitory synaptic conductances of primary auditory cortex neurons. Journal of neurophysiology, 92(1): 630–643, 2004.

[55] Alex M Thomson and Jim Deuchars. Temporal and spatial properties of local circuits in neocortex. Trends in neurosciences, 17(3): 119–126, 1994.

[56] Misha Tsodyks, Klaus Pawelzik, and Henry Markram. Neural networks with dynamic synapses. Neural computation, 10(4):821–835, 1998.

[57] Misha V Tsodyks and Henry Markram. The neural code between neocortical pyramidal neurons depends on neurotransmitter release probability. Proceedings of the national academy of sciences, 94(2):719–723, 1997.

[58] Carl Van Vreeswijk and Haim Sompolinsky. Chaos in neuronal networks with balanced excitatory and inhibitory activity. Science, 274(5293):1724–1726, 1996.

[59] Juan A Varela, Kamal Sen, Jay Gibson, Joshua Fost, LF Abbott, and Sacha B Nelson. A quantitative description of short-term plasticity at excitatory synapses in layer 2/3 of rat primary visual cortex. Journal of Neuroscience, 17(20): 7926–7940, 1997.

[60] Donald A Wilson. Rapid, experience-induced enhancement in odorant discrimination by anterior piriform cortex neurons. Journal of neurophysiology, 90(1):65–72, 2003.

